# A cell atlas of human thymic development defines T cell repertoire formation

**DOI:** 10.1101/2020.01.28.911115

**Authors:** Jong-Eun Park, Rachel A. Botting, Cecilia Domínguez Conde, Dorin-Mirel Popescu, Marieke Lavaert, Daniel J. Kunz, Emily Stephenson, Roberta Ragazzini, Elizabeth Tuck, Anna Wilbrey-Clark, John R. Ferdinand, Simone Webb, Daniel Maunder, Niels Vandamme, Krishnaa Mahbubani, Krzysztof Polanski, Lira Mamanova, Andrew Fuller, Andrew Filby, Gary Reynolds, David Dixon, Kourosh Saeb-Parsy, Steven Lisgo, Deborah Henderson, Roser Vento-Tormo, Kerstin B. Meyer, Yvan Saeys, Paola Bonfanti, Sam Behjati, Menna R. Clatworthy, Tom Taghon, Muzlifah Haniffa, Sarah A. Teichmann

**Affiliations:** Wellcome Sanger Institute, Wellcome Genome Campus, Hinxton, Cambridge CB10 1SA, UK; Institute of Cellular Medicine, Newcastle University, Newcastle upon Tyne, NE2 4HH, UK; Faculty of Medicine and Health Sciences, Department of Diagnostic Sciences, Ghent University, C. Heymanslaan 10, MRB2, Entrance 38, 9000 Ghent, Belgium; Cancer Research Institute Ghent (CRIG), Ghent University, Ghent, Belgium; Molecular Immunity Unit, Department of Medicine, University of Cambridge, MRC Laboratory of Molecular Biology, United Kingdom, CB2 0QQ; Data Mining and Modeling for Biomedicine, VIB Center for Inflammation Research, Ghent, Belgium; Department of Surgery, University of Cambridge and NIHR Cambridge Biomedical Research Centre, United Kingdom, CB2 0QQ; Institute of Genetic Medicine, Newcastle University, Newcastle upon Tyne, NE1 3BZ, UK; The Francis Crick Institute, London, UK; Great Ormond Street Institute of Child Health, University College London, London, UK; Institute of Immunity and Transplantation, University College London, London, UK; Cambridge University Hospitals NHS Foundation Trust, United Kingdom, CB2 0QQ; Department of Dermatology and NIHR Newcastle Biomedical Research Centre, Newcastle Hospitals NHS Foundation Trust, Newcastle upon Tyne NE2 4LP, UK; Theory of Condensed Matter Group, Cavendish Laboratory/Department of Physics, University of Cambridge, Cambridge CB3 0HE, UK; The Wellcome Trust/Cancer Research UK Gurdon Institute, University of Cambridge, Cambridge, United Kingdom; Department of Applied Mathematics, Computer Science and Statistics, Ghent University, Ghent, Belgium; Department of Haematology and Wellcome and MRC Cambridge Stem Cell Institute, University of Cambridge, Cambridge, CB2 2XY, UK; Department of Paediatrics, University of Cambridge, Cambridge CB2 0SP, UK

## Abstract

The thymus provides a nurturing environment for the differentiation and selection of T cells, a process orchestrated by their interaction with multiple thymic cell types. We utilised single-cell RNA-sequencing (scRNA-seq) to create a cell census of the human thymus and to reconstruct T-cell differentiation trajectories and T-cell receptor (TCR) recombination kinetics. Using this approach, we identified and located *in situ* novel CD8αα^+^ T-cell populations, thymic fibroblast subtypes and activated dendritic cell (aDC) states. In addition, we reveal a bias in TCR recombination and selection, which is attributed to genomic position and suggests later commitment of the CD8^+^ T-cell lineage. Taken together, our data provide a comprehensive atlas of the human thymus across the lifespan with new insights into human T-cell development.

## Introduction

The thymus plays an essential role in the establishment of adaptive immunity and central tolerance as it mediates the maturation and selection of T cells. This organ degenerates early during life and the resulting reduction in T-cell output has been linked to age-related incidence of cancer, infection and autoimmunity (*1, 2*). T-cell precursors from fetal liver or bone marrow migrate into the thymus, where they differentiate into diverse types of mature T cells (*3, 4*). The thymic microenvironment, consisting of diverse cell types, cooperatively supports T-cell differentiation (*5, 6*). While thymic epithelial cells (TECs) provide critical cues to promote T-cell fate (*7*), Other cell types are also involved in this process, such as dendritic cells (DC) that undertake antigen presentation, and mesenchymal cells, which support TEC differentiation and maintenance (*8–11*). Seminal experiments in animal models have provided major insights into the function and cellular composition of the thymus (*12, 13*). More recently, scRNA-seq has revealed new aspects of thymus organogenesis and new types of thymic epithelial cells (TECs) in mouse (*14–16*). However, the human organ matures in a mode and tempo that is unique to our species (*17–19*), calling for a comprehensive genome-wide study for human thymus.

T-cell development involves a parallel process of staged T-cell lymphocyte differentiation accompanied by acquisition of a diverse TCR repertoire for antigen recognition (*20*). To achieve the required diversity, TCR sequences are composed of several modules (*i.e.*, V, D and J modules) with multiple genomic copies for each module. Genomic recombination selects and concatenates one copy from each module, with some non-templated modifications added at the junctions. Interestingly, this VDJ gene recombination can preferentially include certain gene segments, leading to the skewing of the repertoire (*21–23*). To date, most of our knowledge of VDJ recombination and repertoire biases, has come from animal models and human peripheral blood analysis, with little comprehensive data on the human thymic TCR repertoire (*22, 24, 25*).

Here, we applied scRNA-seq to generate a comprehensive transcriptomic profile of the diverse cell populations present in embryonic, fetal, paediatric and adult stages of the human thymus. We combined this with detailed TCR repertoire analysis to reconstruct the T-cell differentiation process at unprecedented resolution. This revealed novel insights into the generation of diverse mature lymphoid lineages from thymus, and the transcription factors which orchestrate this process. We provide this data to the scientific community in an interactive web portal (https://developmentcellatlas.ncl.ac.uk).

### Cellular composition of the developing human thymus

We performed scRNA-seq on 11 fetal thymi from 7 post-conception weeks (PCW), when the thymic rudiment can be dissected, to 17 PCW, when thymic development is completed (**Fig. 1, A and B**). We also analysed 4 postnatal samples, covering the entire period of active thymic function. Isolated single cells were FACS-sorted based on CD45 or CD3 expression to separate abundant T lymphocytes from other cell types, and then analysed by single-cell transcriptomics coupled with TCRɑβ profiling. After quality control including doublet removal, we obtained a total of 112,783 cells from the developing thymus and 59,187 cells from postnatal thymus (**Data S1**). If available, other relevant organs (fetal liver as a hematopoietic organ, bone marrow, spleen and lymph nodes) were collected from the same donor. We performed batch correction using the BBKNN algorithm (**fig. S1**) (*26*).

**Fig. 1.**
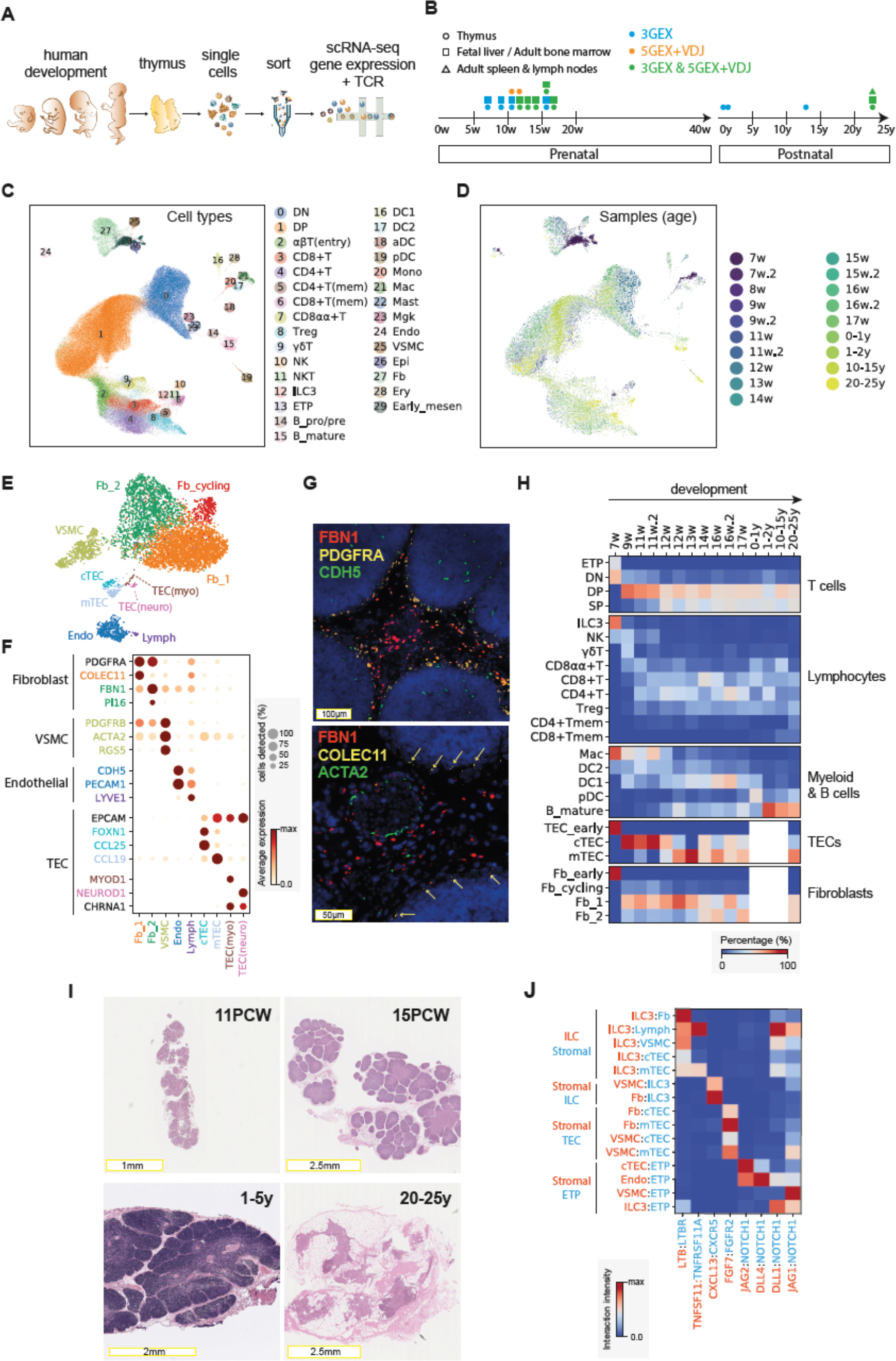
Cellular composition of the developing human thymus. **(A)** Schematic of single-cell transcriptome profiling of the developing human thymus. **(B)** Summary of gestational stage/age of samples, organs (circle: thymus, rectangle: fetal liver and adult bone marrow, triangle: adult spleen and lymph nodes) and 10x Genomics chemistry (colours). **(C)** UMAP visualisation of the cellular composition of the human thymus colored by cell type (DN: double-negative T cells, DP: double-positive T cells, ETP: Early thymic progenitor, B_pro/pre: pro/pre B cells, aDC: activated dendritic cells, pDC: plasmacytoid dendritic cells, Mono: monocyte, Mac: macrophage, Mgk: megakaryocyte, Endo: endothelial cells, VSMC: vesicular smooth muscle cells, Epi: epithelial cells, Fb: fibroblasts, Ery: erythrocytes, Early_mesen: early mesenchymal cells). (**D)** Same UMAP plot coloured by age, indicated by post-conception weeks (PCW) or postnatal years (y). Samples are colour coded based on age groups. **(E)** UMAP visualisation of thymic epithelial, endothelial and fibroblast cell types and **(F)** dot plot of their marker genes. Color represents maximum-normalised mean expression of marker genes in each cell group, and size indicates the proportion of cells expressing marker gene. (This scheme is consistently used throughout the manuscript.) **(G)** RNA single-molecule FISH in 15 PCW thymus slide with probes targeting stromal cell populations. Top panel: Fb2 population marker *FBN1* (red), general fibroblast markers *PDGFRA* (yellow) and *CDH5* (green). Lower panel: Fb2 marker *FBN1* (red), Fb1 markers *COLEC11* (yellow) and *ACTA2* (green). Data representative of n=2. **(H)** Relative abundance of cell types with age, color represents percentage within each cellular subgroup. **(I)** H&E staining of cross-sectioned thymic tissue at different developmental and postnatal ages. **(J)** Predicted cell-cell interactions showing receptor-ligand signaling mediating thymic development. Color represents intensity of interaction calculated by multiplying the mean expression of receptor-ligand pairs and normalising to maximum per interaction. (This scheme is consistently used throughout the manuscript).

We have annotated cell clusters into at least 30 different cell types or cell states (**Fig. 1, C and D**), which can be clearly identified by the expression of specific marker genes (**fig. S2**). Differentiating T cells are well represented in the dataset, including double negative (DN), double positive (DP), CD4^+^ single positive (CD4^+^T), CD8^+^ single positive (CD8^+^T), FOXP3^+^ regulatory (Treg), CD8αα^+^ and γδ T cells. We also identified other immune cells including B cells, NK cells, innate lymphoid cells (ILCs), macrophages, monocytes and dendritic cells (DCs).

Our dataset also featured diverse non-immune cell types, which constitute the thymic microenvironment. We further classified them into subtypes including fibroblasts, vascular smooth muscle cells (VSMCs), endothelial cells, lymphatic endothelial cells and thymic epithelial cells (TECs) (**Fig. 1, E and F**).

Thymic fibroblasts were further divided into two subtypes, neither of which has been previously described: Fibroblast type 1 (Fb1) cells (*COLEC11*, *C7, GDF10*) and Fibroblast type 2 (Fb2) cells (*PI16, FN1*, *FBN1)* (**Fig. 1F**). Fb1 cells uniquely express *ALDH1A2*, an enzyme responsible for the production of retinoic acid, which regulates epithelial growth (*27*). In contrast, extracellular matrix (ECM) genes and Semaphorins (*SEMA3C*, *SEMA3D*), which regulate vascular development (*28*), are specifically detected in Fb2 (**fig. S3A**). To explore the localisation pattern of fibroblast subtypes, we performed *in situ* smFISH targeting Fb1 and Fb2 markers (*COLEC11* and *FBN1*) together with general fibroblast (*PDGFRA*), endothelial (*CDH5*) and VSMC (*ACTA2*) markers (**Fig. 1G**). The results show that Fb1 cells were peri-lobular, while Fb2 cells were interlobular, often associated with large blood vessel lined with VSMCs, consistent with their transcriptomic profile of genes regulating vascular development.

In addition to fibroblasts, we also identified subpopulations of human thymic epithelial cells, which have not been characterised before with genome-wide profiling (**Fig. 1E**). The majority of the epithelial cells in our data correspond to the known cortical and medullary TECs (cTECs and mTECs), identified by canonical marker genes such as *CCL25*, *KRT5* for cTECs, and *CCL19*, *KRT8*, *AIRE* for mTECs (**Fig. 1F**). *FOXN1*, the master transcriptional regulator for TEC development, was more highly expressed in cTECs. In addition to these conventional TECs, two other less well known EPCAM^+^ cell types were discernible: *MYOD1* and *MYOG*-expressing myoid cells (TEC(myo)s) and *NEUROD1*, *SYP*, *CHGA*-expressing TEC(neuro)s (**Fig. 1F**). Notably, *CHRNA1*, which has been associated with the autoimmune disease myasthenia gravis (*29*), was specifically expressed by both of these cell types, supporting the notion that antigen expression by TEC(myo) and TEC(neuro) cells may be involved in tolerance induction (*30, 31*).

Lastly, we analysed the expression pattern of genes known to cause congenital T-cell immunodeficiency to provide insight into when and where these rare disease genes may play a role during thymic development (**fig. S4**). We find three major patterns: (1) genes highly expressed in T cells, involved in T-cell recombination, signalling and migration; (2) genes broadly expressed in immune cells, involved in cell migration, antigen processing and MHC transcription; and (3) genes highly expressed in the stromal and epithelial compartment, which underlie structural defects of the thymus. Interestingly, we found specific expression of CHARGE syndrome gene *SEMA3E* in epithelial cells, emphasizing the role of epithelial cells in thymocyte trafficking (*32*).

### Coordinated development of thymic stroma and T cells

Next, we investigated the dynamics of relative proportions of different cell types across development (**Fig. 1H**). In the earliest fetal sample (7 PCW), the lymphoid compartment contained NK and ILC3s, with very few differentiating αβT cells. Moreover, stromal cells were not yet fully differentiated, with progenitor states present in both the fibroblast and epithelial cell compartments (**Fig. 1H** and **fig. S5, A and B**).

From 9 PCW onwards, increasing numbers of DN and DP T cells were observed. The fraction of single-positive mature T cells (SP) gradually increased thereafter, reaching equilibrium at around 12 PCW (**Fig. 1H**). Conversely, the proportion of innate lymphocytes decreased (**Fig. 1H**).

Of note, more diverse mature T-cell types were present in the young adult sample, which showed evidence of degeneration in thymic morphology (**Fig. 1I**). Comparison with spleen and lymph nodes taken from the same donor showed the presence of terminally differentiated T cells in the thymus, suggesting re-entry into thymus or contamination with circulating cells (**Fig. 1H and fig. S5, C and D**). Notably, cytotoxic CD4^+^T lymphocytes (CD4^+^CTL) expressing *IL10*, perforin and granzymes were enriched in the degenerated thymus sample, implicating a potential role for these cells in thymic involution (*33*) (**fig. S5E**).

The trend in T cell development was mirrored by corresponding changes in thymic stromal cells. We observed temporal changes in TEC populations from the early, precursor TEC population towards differentiated cTECs and mTECs (**Fig. 1H**), aligned with the onset of T-cell maturation. This supports the notion of ‘thymic crosstalk’ in which epithelial cells and mature T cells interact synergistically to support their mutual differentiation (*34*). The progression in cellular development was reflected in gross tissue morphology, showing the formation of the medulla only after 11 PCW (**Fig. 1I**).

Moreover, fibroblast composition also changed during development. The Fb1 population mentioned in the previous section dominated in early development, with similar numbers of Fb1 and Fb2 cells observed at later developmental timepoints, and a reduction in the number of cycling cells (**Fig. 1H**). This reflects an early expansion of epithelial lobules followed by maturation of the connective tissue. We validated thymic fibroblast explant cultures, and show the increase in PI16 protein level, a marker for Fb2 cells, by FACS analysis (**figs. S3, B and C**).

Finally, other immune cells also change dynamically over gestation and in postnatal life. Macrophages were abundant at early stages, while DCs and B cells increased throughout development (**Fig. 1H**). Among DCs, DC2 predominated in early development, followed by the appearance of DC1 after 12 PCW, and pDC only in postnatal life.

To further investigate the factors mediating the coordinated development of thymic stroma and T cells, we systematically investigated cellular interactions using our public database CellPhoneDB (*35*) to predict the ligand-receptor pairs between them. (**Data S2**). This showed lymphotoxin signaling (*LTB*:*LTBR*) from ILC3 to diverse stromal cells, and RANKL-RANK (*TNFRSF11*:*TNFRSF11A*) from ILC3 to mTECs and lymphatic endothelial cells (**Fig. 1J**). Furthermore, the predicted results suggest that stromal cells potentially recruit ILC3 by *CXCL13*:*CXCR5* interaction, and induce TEC development by *FGF7*:*FGFR2* signalling (**Fig. 1J**). In addition, we analysed the expression pattern of Notch ligands and receptors, responsible for T-cell commitment. While *NOTCH1* was the main receptor expressed in ETPs, we observed that multiple cells expressed different Notch ligands: cTECs and endothelial cells expressed both *JAG2* and *DLL4*, VSMCs expressed *JAG1*, and ILC3 expressed *DLL1* and *JAG1* (**Fig. 1J**) (*36*). Taken together, our dataset demonstrates the interplay between diverse cell types, which mediate coordinated development of the thymus.

### Conventional T cell differentiation trajectory

As fetal liver is the main haematopoietic organ and source of HSC/MPP when the thymic rudiment develops, we analysed paired thymus and liver samples from the same fetus. This allowed us to investigate the relationship between fetal liver hematopoietic stem cells/multipotent progenitors (HSC/MPP) and thymic progenitors. We merged the thymus and liver data, and selected clusters including liver HSC/MPP, thymic ETPs and DN thymocytes for data analysis and visualisation (**Fig. 2, A and B**). This positioned thymic ETPs at the isthmus between fetal liver HSC/MPP and pre/pro B cells. Interestingly, progenitors from 7 PCW thymus (**Fig. 2B**) showed a highly skewed transcriptomic profile adjacent to that of B and T cells, compared to fetal liver progenitors from the same donor (**Fig. 2B**). This observation was also made at later developmental stages, when T cell differentiation is evident as a peninsula extending from the ETP and DN clusters (**Fig. 2B, orange**). Our analysis demonstrates that fetal thymic seeding populations are skewed towards lymphoid potential.

**Figure 2.**
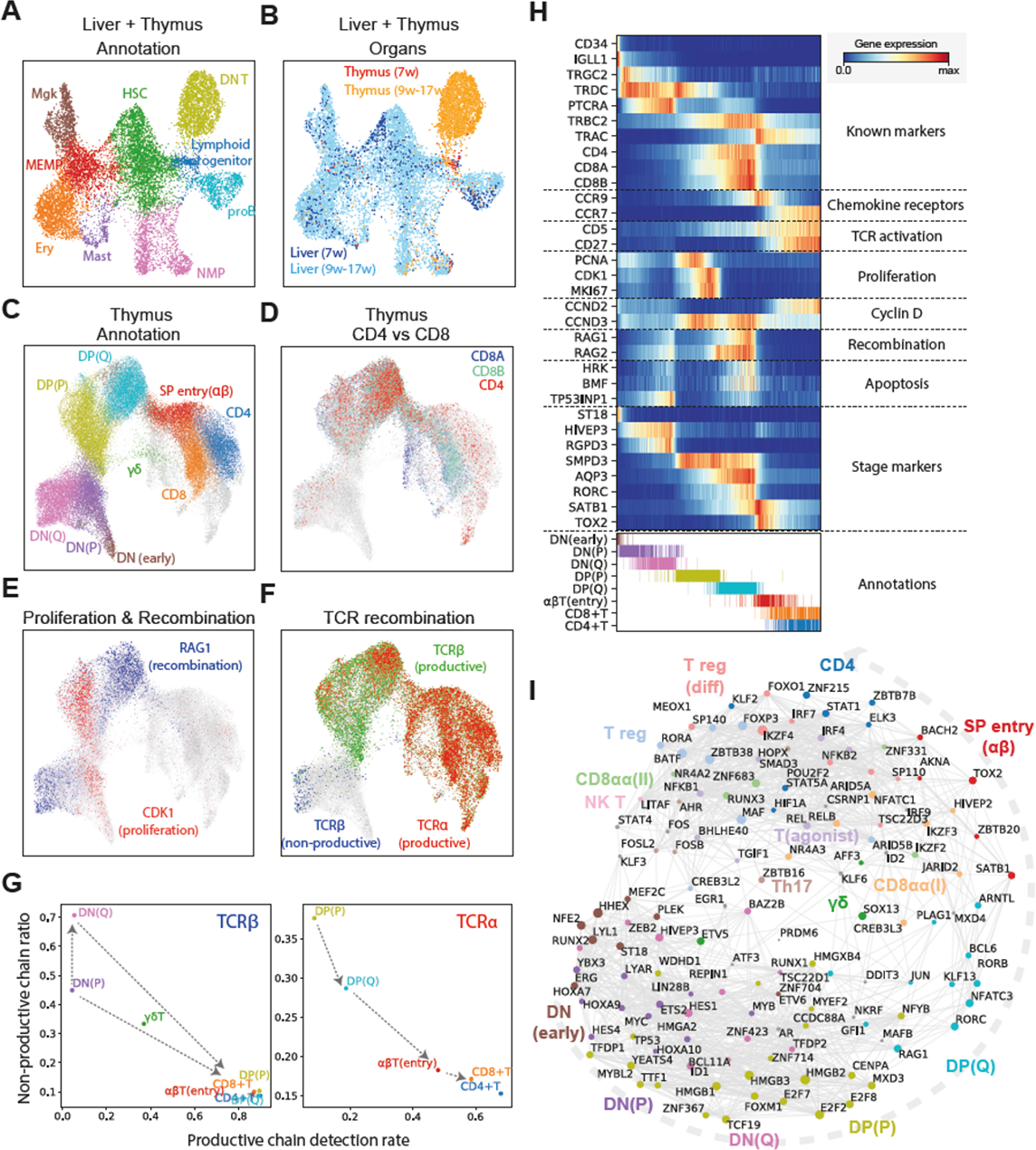
Thymic seeding of early thymic progenitors (ETPs) and T cell differentiation trajectory. **(A)** UMAP visualisation of ETP and fetal liver hematopoietic stem cells/early progenitors. (HSC: Hematopoietic stem cells, ETP: Early T cell progenitors, DN: Double negative T cells, NMP: Neutrophil-Myeloid progenitors, MEMP: Megakaryocyte-Erythrocyte-Mast cell progenitors, Ery: Erythrocytes, Mgk: Megakaryocytes). The same UMAP coloured by **(B)** organ (liver in blue and thymus in yellow/red). **(C)** UMAP visualisation of developing thymocytes after batch correction. (DN: double negative T cells, DP: double positive T cells, SP: single positive T cells, P: proliferating, Q: quiescent). The data contains cells from all sampled developmental stages. Cells from abundant clusters are down-sampled for better visualisation. The reproducibility of structure is confirmed across individual sample. Unconventional T cells are marked as grey. (**D-F**) The same UMAP plot showing *CD4* (red), *CD8A* (blue) and *CD8B* (turquoise) gene expression (D), *CDK1* (red) cell cycle and *RAG1* (blue) recombination gene expression (E), and TCRα (red) and TCRβ (green = productive and blue = non-productive) VDJ genes (F). **(G)** Scatter plot showing the rate of productive chain detection within cells in specific cell types (x-axis) and the ratio between the number of non-productive/productive TCR chains detected in specific cell types (y-axis); TCRβ (left panel) and TCRα (right panel). **(H)** Heatmap showing differentially expressed genes across T cell differentiation pseudotime from T DN to CD4+ and CD8+ T cells. Upper panel: X-axis represents pseudo-temporal ordering. Gene expression levels across pseudotime axis are maximum-normalised and smoothed. Genes are grouped by their functional categories and expression patterns. Lower panel: Cell type annotation of cells aligned along the pseudotime axis. The same colour schemes are used as (C). **(I)** Graph showing correlation-based network of transcription factors expressed by thymocytes. Nodes represent transcription factors, and edge widths are proportional to the correlation coefficient between two transcription factors. TFs with significant association to specific cell types depicted in colour. Node size is proportional to the significance of association to specific cell types.

To investigate the downstream T cell differentiation trajectory, we selected the T cell populations and projected them using UMAP and force-directed graph analysis (**Fig. 2C, fig. S6A and Data S3),** which showed a continuous trajectory of differentiating T cells. To confirm the validity of this trajectory, we overlaid hallmark genes of T-cell differentiation: CD4/CD8A/CD8B genes (**Fig. 2D**), cell cycle (*CDK1*) and recombination (*RAG1*) genes (**Fig. 2E**) and fully recombined TCRɑ/TCRβ (**Fig. 2F**). The trajectory started from CD4^-^CD8^-^ DN cells, which gradually express CD4 and CD8 to become CD4^+^CD8^+^ DP cells, and then diverge into mature CD4^+^ or CD8^+^ SP cells (**Fig. 2D**). We also noted a separate lineage of cells diverging from the DN-DP junction corresponding to γδ T-cell differentiation. Additional T-cell lineages identified in this analysis will be discussed in the following section (**Fig. 2C, grey**).

Notably, DN and DP cells were separated into two phases by the expression of cell cycle genes (**Fig. 2E**). We designated the early population with strong cell cycle signature as proliferating (P) and the later population quiescent (Q), respectively (**Fig. 2C**). Expression of VDJ recombination genes (*RAG1* and *RAG2*) increased from the late proliferative phase, and peaked at the quiescent phases. This pattern reflects the proliferation of T cells which precedes each round of recombination.

Next, we aligned the TCR recombination data to this trajectory (**Fig. 2F**). In the DN stage, recombined TCRβ sequences were detected from the late P phase, which coincided with the increase in recombination signature. The ratio of non-productive to productive recombination events (non-productivity score) for TCRβ was relatively higher in DN stages, and dropped to a basal level as cells entered DP stages, demonstrating the impact of beta-selection (**Fig. 2G**).

Notably, the non-productivity score for TCRβ was highest in the DN(Q) stage, suggesting that cells failing to secure a productive TCRβ recombination for the first allele undergo recombination of the other allele. In the DP stage, recombined TCRɑ chains were detected from P stage onwards. In contrast to TCRβ, non-productive TCRɑ chains were not enriched in the DP(Q) cells, but were rather depleted (**Fig. 2G**). These data reflect differences between the kinetics of beta-selection and alpha-selection, which will be further discussed later.

To model the development of conventional ɑβT cells in more detail, we performed pseudo-time analysis, which resulted in an ordering of cells highly consistent with known marker genes and transcription factors (**Fig. 2H**). In addition, we identified novel T-cell developmental markers, including *ST18* for early DN, *AQP3* for DP and *TOX2* for DP to SP transition. To derive further insights into transcription factors that specify T-cell stages and lineages, we created a correlation-based transcription factor network, after imputing gene expression (see Methods). The resulting network demonstrated modules of transcription factors specific to each stage of T-cell differentiation and lineage commitment (**Fig. 2I**).

### Development of Tregs and discovery of *GNG4*^+^ CD8ɑɑ T cells in the thymic medulla

In addition to conventional CD4^+^ or CD8^+^ T cells, which comprise the majority of T cells in the developing thymus, our data identified multiple unconventional T cell types, which were grouped by the expression of signature marker genes (**Fig. 3, A, B** and **Fig. 2I**). Unconventional T cells have been suggested to require agonist selection for development (*3*). In support of this, we observed a lower ratio of non-productive TCR chains for these cells, implying that they reside longer in the thymus compared to conventional T cells (**Fig. 3C**).

**Figure 3.**
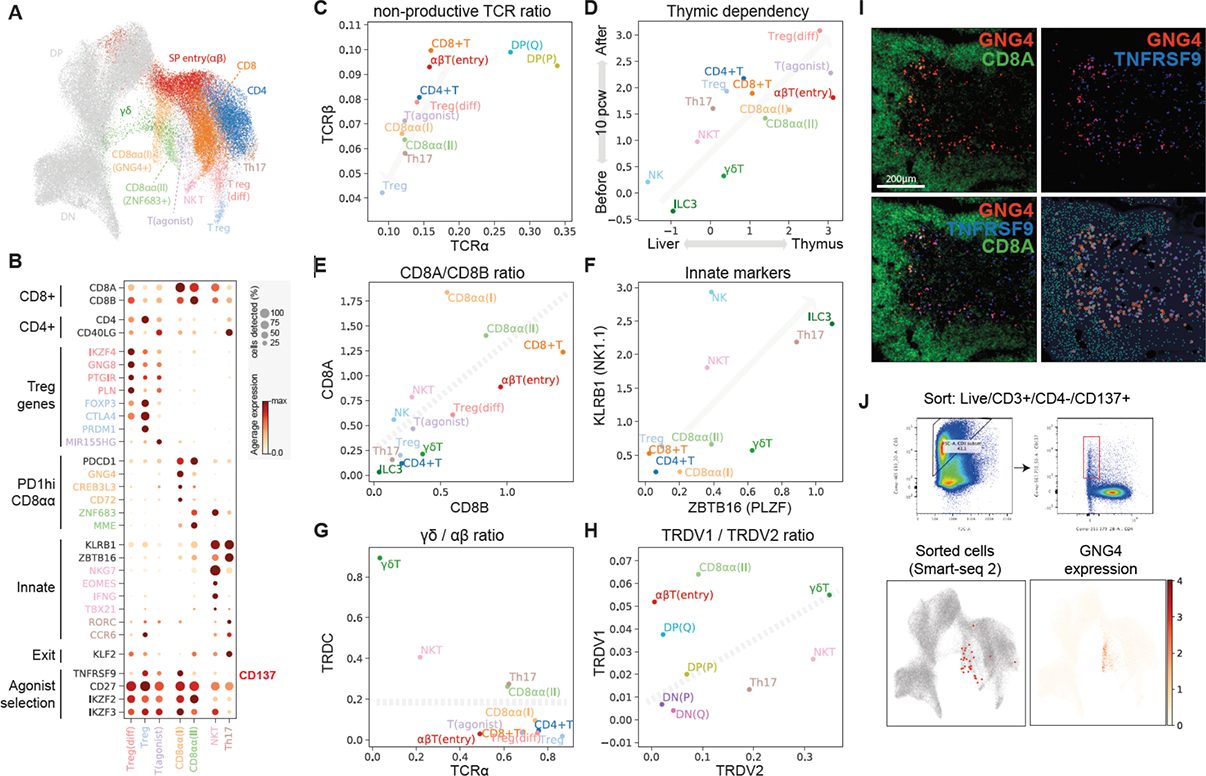
Identification of GNG4+ CD8aa T cells in the thymic medulla. **(A)** UMAP visualisation of mature T cell populations in the thymus. Axes and coordinates are as Fig. 2C. (The cell annotation colour scheme used here is maintained throughout this figure.) **(B)** Dot plot showing marker gene expression for the mature T cell types. Genes are stratified according to associated cell types or functional relationship. **(C)** Scatter plot showing the ratio between the number of non-productive/productive TCR chains detected in specific cell types in TCRα chain (x-axis) and TCRβ chain (y-axis). Same colour schemes apply as in (A). The grey arrow indicates a trendline for decreasing non-productive TCR chain ratio in unconventional *versus* conventional T cells. **(D)** Scatter plot showing the relative abundance of each cell type between fetal liver and thymus (x-axis) and before and after thymic maturation (delimited at 10 PCW) (y-axis). Grey arrow indicates trendline for increasing thymic dependency. **(E-H)** Scatter plot comparing the characteristics of unconventional T cells based on *CD8A* vs. *CD8B* expression levels (E), *KLRB1* vs *ZBTB16* expression levels (F), TCRα productive chain vs *TRDC* detection ratio (G) and *TRDV1* vs *TRDV2* expression levels (H). Grey arrows or lines are used to set boundaries between groups (E, G, H) or indicate the trend of innate marker gene expression (F). **(I)** single-molecule RNA FISH showing *GNG4* (red), *TNFRSF9* (blue) and *CD8A* (green) in a 15 PCW thymus. Right bottom panel shows detected spots from the image on top of the tissue structure based on DAPI signal. Colour scheme for spots are the same as in the image. **(J)** FACS gating strategy to isolate CD8aa(I) cells (live/CD3+/CD4-/CD137+) and Smart-seq2 validation of FACS-isolated cells projected to the UMAP presentation of total mature T cells from discovery dataset (bottom left panel). *GNG4* expression pattern is overlayed onto the same UMAP plot (bottom right panel).

Next, we investigated whether development of these unconventional T cells was dependent on the thymus. We reasoned that if a population is thymus-dependent, it would accumulate after thymic maturation (∼10 PCW) and be enriched in the thymus compared to other hematopoietic organs. Consistent with this, all unconventional T cells were enriched in the thymus, particularly post-thymic maturation, suggesting that they are thymus-derived (**Fig. 3D**).

Tregs were the most abundant unconventional T cells in the thymus. There was a clear differentiation trajectory connecting ɑβT cells and Tregs. We defined the connecting population as differentiating Tregs (Treg(diff)) (**Fig. 3A)**. Compared to canonical Tregs, Treg(diff) cells had lower *FOXP3* and *IL2RA* expression, and higher expression of *IKZF4*, *GNG8*, *PTGIR* and *PLN* (**Fig. 3B)**. Of note, *IKZF4*, *GNG8* and *PTGIR* have been associated with autoimmunity and Treg differentiation (*37*). This highlights the relevance of the genes identified by our analysis and underscoring the importance of the intermediate cell states during differentiation for avoidance of autoimmunity.

Lastly, we noted another intermediate population which shares expression modules with Treg(diff) cells, but not with terminally differentiated Treg cells. We named this population as T(agonist) and found a specific marker non-coding RNA, MIR155HG (**Fig. 3, A and B)**.

Other unconventional T cell populations included CD8ɑɑ^+^T cells, NKT-like cells and Th17-like cells (**Fig. 3B**). There were three distinct populations of CD8ɑɑ^+^T cells: *GNG4*^+^ *CD8ɑɑ*^+^T(I) cells, *ZNF683*^+^ *CD8ɑɑ*^+^T(II) and a *CD8ɑɑ*^+^ NKT-like population marked by *EOMES* (**Fig. 3E**). *GNG4*^+^*CD8ɑɑ*^+^T(I) and *ZNF683*^+^*CD8ɑɑ*^+^T(II) both shared *PDCD1* expression at an early stage, which decreased in their terminally differentiated state (**fig. S6B**). While *GNG4*^+^ *CD8ɑɑ*^+^T(I) displayed a clear trajectory diverging from late DP stage (ɑβT SP entry cells), *ZNF683*^+^*CD8ɑɑ*^+^T(II) cells have a mixed ɑβ and γδ T cell signatures, and sit next to both *GNG4*^+^*CD8ɑɑ*^+^T(I) cells and γδ T cells (**Fig. 3A and fig. S6B**).

*EOMES*^+^ NKT-like cells have a shared gene expression profile with NK cells (*NKG7*, *IFNG*, *TBX21*) and are enriched in γδ T cells, i.e. their TCRs are γδ rather than ɑβ (**Fig. 3B and fig. S6**). Interestingly, previously described gene sets from bulk RNA-sequencing of human thymic or cord blood CD8ɑɑ^+^T cells can now be deconvoluted into our three CD8ɑɑ^+^T cell populations using signature genes. These results suggest that our three novel CD8ɑɑ^+^T cell populations are present in these previously published thymic and cord blood samples at different frequencies, as shown in (**fig. S7**) (*38*).

Finally, we found another fetal specific cell cluster which we named as “Th17-like cells”, based on *CD4*, *CD40LG*, *RORC* and *CCR6* expression (**Fig. 3B**). Th17-like cells and NKT-like cells expressed *KLRB1* and *ZBTB16*, which are hallmarks of T cells with innate character (*39, 40*) (**Fig. 3F**).

As described above, many cell clusters contained a mixed signature of ɑβ and γδ T cells, meaning that a single cluster contained some cells with ɑβ TCR expression and others with γδ TCR. To classify cells into ɑβ and γδ T cells, we analysed the TCRɑ/δ loci, where recombination of TCRɑ excises TCRδ, making the two mutually exclusive (**Fig. 3G)**. This clearly showed that γδ T cells diverging between the DN and DP populations are pure γδ T cells. In contrast, CD8ɑɑ^+^T(II), NKT-like and Th17-like cells include both ɑβ and γδ T cell populations, suggesting transcriptomic convergence of some ɑβ and γδ T cells.

Interestingly, *TRDV1* and *TRDV2*, the two most frequently used TCRδ V genes in human, displayed clear usage bias: *TRDV2* was used at an earlier stage (DN), while *TRDV1* was exclusively utilised in later T-cell development (DP(Q) and ɑβT entry) (**Fig. 3H**). Based on this pattern, we can attribute the stage of origin of γδ T-cell populations, which suggests that CD8ɑɑ^+^T(II) are derived from the late DP stage, while NKT-like/Th17-like cells arise from earlier stages (**Fig. 3H**).

Having identified unconventional T cells and their trajectory of origin within thymic T-cell development, we focused on our newly discovered GNG4^+^CD8ɑɑ^+^T(I) cells, as they have a unique gene expression profile (*GNG4*, *CREB3L3* and *CD72*). This is in contrast to CD8ɑɑ^+^T(II) cells, which express known markers of CD8ɑɑ^+^T cells such as *ZNF683* and *MME* (*38*). Moreover, the expression level of *KLF2*, a regulator of thymic emigration, was extremely low in CD8ɑɑ^+^T(I) cells, suggesting that they may be thymic-resident (**Fig. 3B**). To locate and validate CD8ɑɑ^+^T(I) cells *in situ*, we performed RNA smFISH targeting *GNG4* in fetal thymus tissue sections. The *GNG4* RNA probe identified a distinct group of cells enriched in the thymic medulla, and co-localised with *CD8A* RNA (**Fig. 3I**). *TNFRSF9* (CD137), is a marker shared between CD8ɑɑ^+^T(I) cells and Tregs. When tested *in situ*, GNG4^+^ cells were a subset of TNFRSF9^+^ cells, further confirming the validity of the localisation pattern.

As CD137 is a surface marker of both CD8ɑɑ^+^T(I) cells and Tregs, we enriched these cells using this marker (**fig. S8**). Further refinement using CD3^+^CD137^+^CD4^-^ FACS-sorting allowed us to specifically enrich for CD8ɑɑ^+^T(I) cells, and confirm their identity by Smart-seq2 scRNA sequencing, providing very deep transcriptomic phenotyping of these cells (**Fig. 3J**).

### Recruitment and activation of DCs for thymocyte selection

Selection of T cells is coordinated by specialised TECs and DCs. We identified three previously well-characterised thymic DC subtypes: DC1 (XCR1^+^CLEC9A^+^), DC2 (SIRPA^+^CLEC10A^+^), pDC (IL3RA^+^CLEC4C^+^) (*6, 41, 42*). We also identified a population that was previously only incompletely described by heterogenous nomenclature, which we term as “activated DCs” (aDCs) (LAMP3^+^CCR7^+^) (**Fig. 4, A and B**) (*43, 44*). aDCs expressed high level of chemokines and co-stimulatory molecules, together with transcription factors like *AIRE* and *FOXD4*, which we validated *in situ* for subset of cells (**Fig. 4B and fig. S9**), suggesting that they may include the previously described AIRE^+^CCR7^+^ DCs in human tonsils and thymus (*45*).

**Figure 4.**
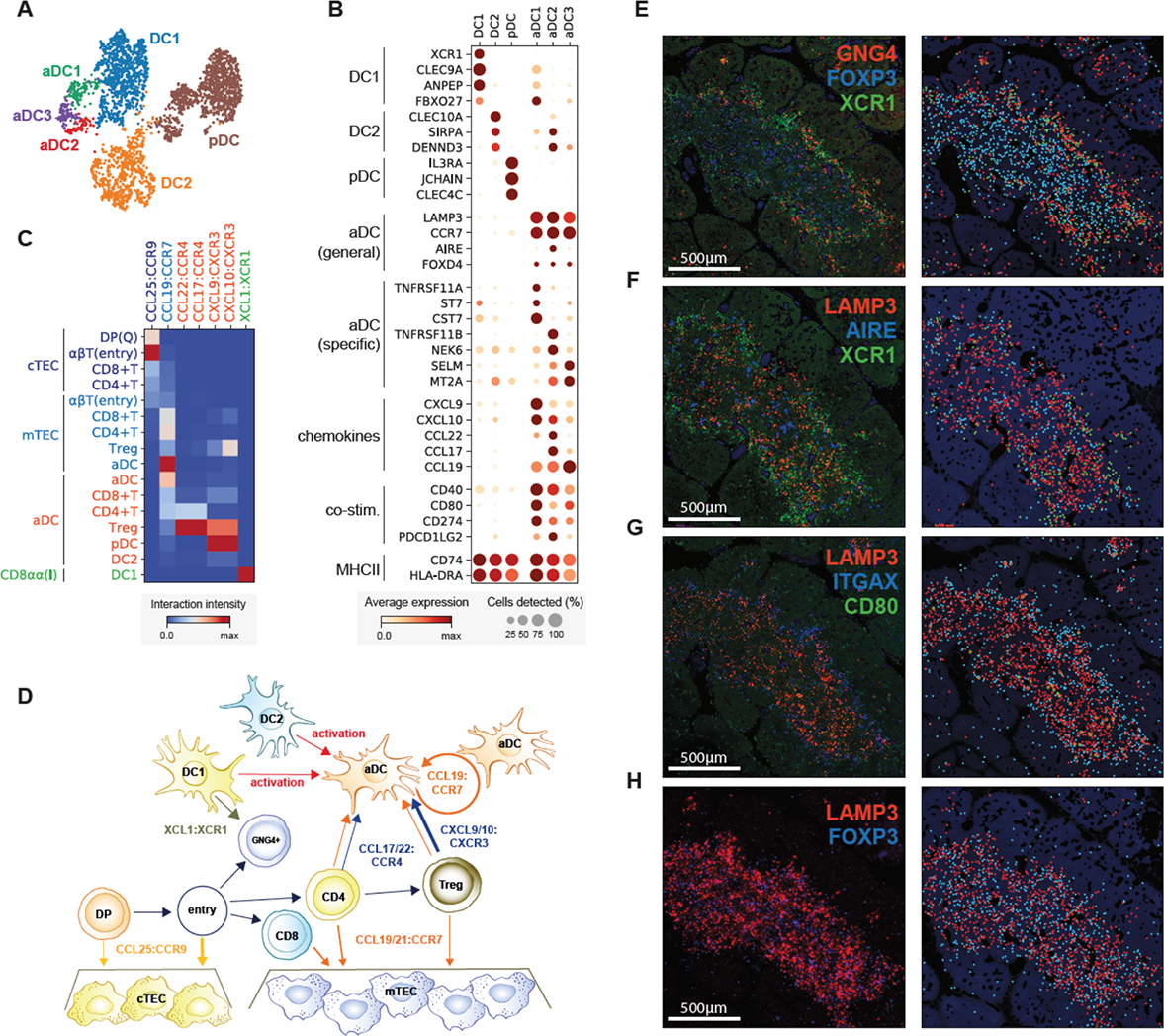
Recruitment and activation of dendritic cells for thymocyte selection. **(A)** UMAP visualisation of thymic DC populations and **(B)** dot plot of their marker genes. **(C)** Heat map of chemokine interactions between T cells, DCs and TECs, where the chemokine is expressed by the outside cell type and the cognate receptor by the inside cell type. **(D)** Schematic model summarising the interactions between thymic epithelial cells (TECs), dendritic cells (DCs) and T cells. The ligand is secreted by the cell at the beginning of the arrow, and the receptor is expressed by the cell at the end of the arrow. **(E)** Left-hand panels: single molecule RNA FISH detection of *GNG4* (red), *XCR1* (green) and *FOXP3* (blue) in 15 PCW thymus. Right-hand panels: Computationally detected spots are presented as a solid circle over the tissue structure based on DAPI signal. Colour schemes for circles are the same as in the image. **(F-H)** Sequential slide sections from the same sample are stained for the detection of *LAMP3* (red), *AIRE* (blue) and *XCR1* (green) (F), *LAMP3* (red), *ITGAX* (blue) and *CD80* (green) (G), *LAMP3* (red), *FOXP3* (blue) (H). Spot detection and representation as in (**E**). Data representative of n = 2.

Interestingly, our single-cell data revealed three subsets within the aDC group, identified by distinct gene expression profiles: aDC1, aDC2 and aDC3 (**Fig. 4, A and B**). aDC1 and aDC2 subtypes shared several marker genes with DC1 and DC2, respectively. To systematically compare aDC subtypes to canonical DCs, we calculated an identity score for each DC population by summarising marker gene expression. This demonstrated a clear relationship between aDC1-DC1 and aDC2-DC2 pairs, suggesting that each aDC subtype derives from a distinct DC population (**fig S10**). Interestingly, aDC1 and aDC2 displayed distinct patterns of chemokine expression, suggesting functional diversification of these aDCs (**Fig. 4B**). Moreover, aDC3 cells had decreased MHC class II and co-stimulatory molecule expression compared to other aDC subsets, which may reflect a post-activation dendritic cell state.

Having identified two canonical TECs and a variety of DC subsets, we used CellPhoneDB analysis to identify specific interactions between these antigen-presenting cells and differentiating T cells (*35*). We focused on interactions mediated by chemokines, which enable cell migration and anatomical co-localisation (**Fig. 4C**). This demonstrated the relay of differentiating T cells from the cortex to the medulla, which is orchestrated by *CCL25*:*CCR9* and *CCL19/21*:*CCR7* interactions between cTEC/mTEC and DP/SP T cells, respectively (*46*).

Interestingly, aDC expressed *CCR7*, together with *CCL19*, enabling attraction to and recruitment of T cells into the thymic medulla. Moreover, they strongly expressed the chemokines *CCL17* and *CCL22*, whose receptor *CCR4* was enriched in CD4^+^ T cells and particularly Tregs. aDCs also potentially recruit other DCs and mature Tregs via *CXCL9/10*:*CXCR3* interactions and are able to provide a strong co-stimulatory signal, which suggests a role in Treg generation. We also noted that GNG4^+^CD8ɑɑ^+^T(I) T cells expressed XCL1, which is involved in the recruitment of XCR1-expressing DC1 cells.

In contrast to the mouse thymus, where XCL1 is mainly expressed by mTECs (*47*), our analysis shows that XCL1 is expressed most highly by CD8ɑɑ^+^T(I) cells and at a lower level by NK cells (**fig. S11**). The location of CD8ɑɑ^+^T(I) in the peri-medullary region suggests a relay of signals from CD8ɑɑ^+^T(I) to recruit XCR1^+^DC1s into the medulla, where these cells are activated and upregulate CCR7. This then leads to migration into the center of the thymic medulla where thymic selection takes place (**Fig. 4D**).

To confirm our *in-silico* predictions, we performed smFISH to identify the anatomical location of CD8ɑɑ^+^T(I) cells (*GNG4*), DC1s (*XCR1*), aDCs (*LAMP3*, *CD80*) and Tregs (*FOXP3*). A generic marker of non-activated DCs (*ITGAX*) and mTECs (*AIRE*) were also used to provide a reference for the organ structure. Imaging of consecutive sections of fetal thymus (15 PCW) revealed the zonation of CD8ɑɑ^+^T(I)/DC1/non-activated DCs located in the peri-medullary region and aDC/Tregs enriched in the center of the medulla (**Fig. 4E-4H**). This is consistent with our model, demonstrating the power of single-cell transcriptomic analysis. Previous studies in mouse models have shown the role of DC and *XCL1* in Treg generation (*43, 47, 48*), suggesting that the signal relay between CD8ɑɑ^+^T(I)/DC1/aDCs may represent an important axis for thymic selection.

### Bias in human TCR repertoire formation and selection

As our data featured detailed T-cell trajectories combined with single-cell resolution TCR sequences, it provided an opportunity to investigate the kinetics of TCR recombination. TCR chains detected from the TCR-enriched 5’ sequencing libraries were filtered for full-length recombinants, and were associated with our cell type annotation. This allowed us to analyse patterns in TCR repertoire formation and selection (**Fig. 5, A and B**).

**Figure 5.**
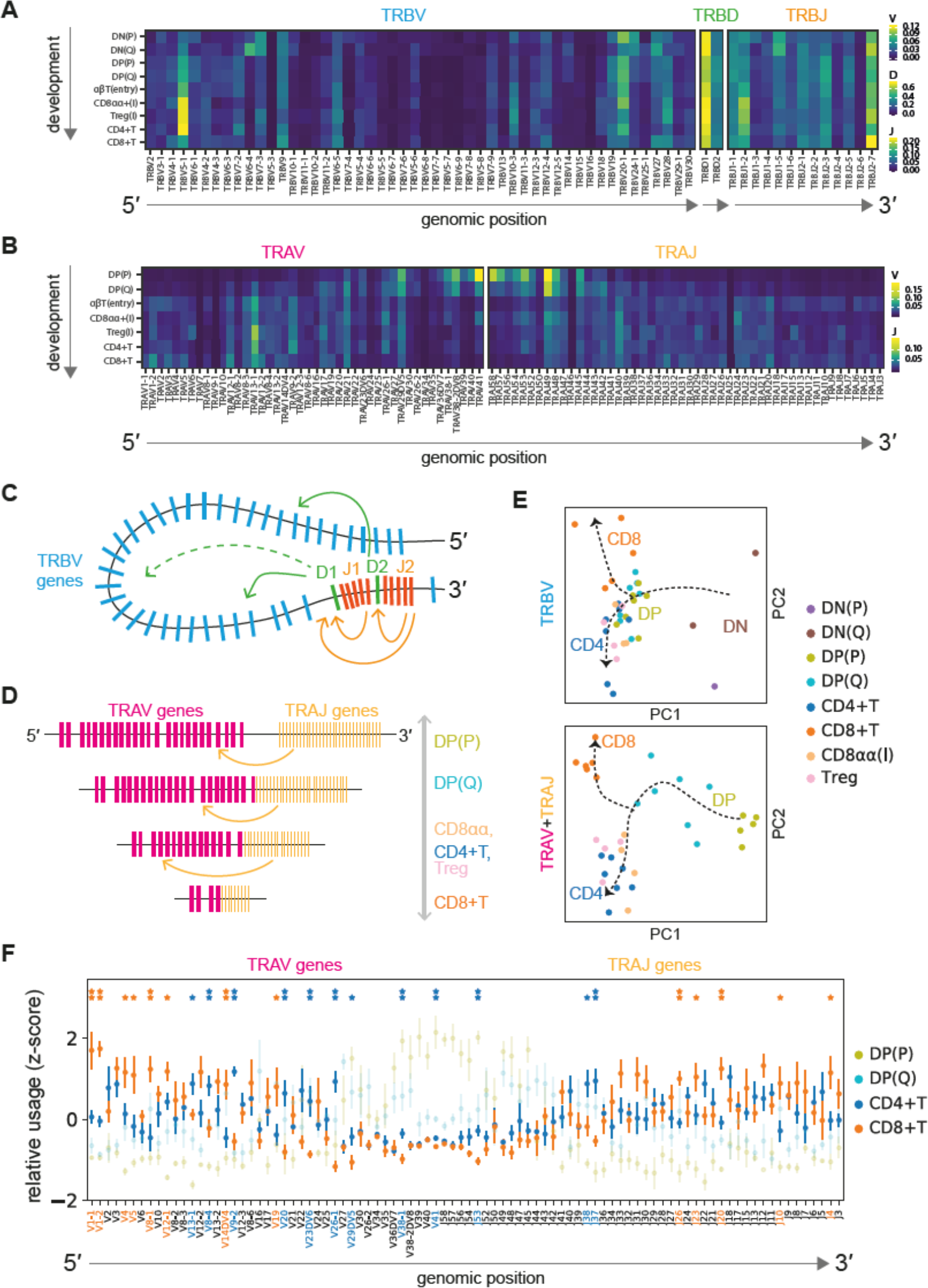
Intrinsic bias in human TCR repertoire formation and selection. **(A)** Heatmap showing the proportion of each TCRβ V, D, J gene segment present at progressive stages of T cell development. Gene segments are positioned according to genomic location. Proportions are calculated from pooled embryonic and fetal data. (The corresponding adult data analysis is presented in fig. S12.) There is consistent enrichment of 5’ and 3’ V segment usage compared to segments in the middle of the locus throughout development. **(B)** Same scheme as in (A) applied to TCRα V and J gene segments. While there is a usage bias of segments at the beginning of development, segments are evenly used by the late developmental stages, indicating progressive recombination leading to even usage of segments. **(C-D)** Schematics illustrating a hypothetical chromatin loop that may explain genomic location bias in recombination of TCRβ locus (C) and the mechanism of progressive recombination of TCRα locus leading to even usage of segments (D). **(E)** PCA plots showing TRBV or TRAV and TRAJ gene usage pattern in different T cell types. Arrows depict T cell developmental order. For TRBV, there is a strong effect from beta selection, after which point the CD4+ and CD8+ repertoires diverge. The development for TRAV+TRAJ is more progressive, with stepwise divergence into the CD4+ and CD8+ repertoires. **(F)** Relative usage of TCRα V and J gene segments according to cell type. The *Z*-score for each segment is calculated from the distribution of normalised proportions stratified by the cell type and sample. *P*-value is calculated by comparing z-scores in CD4+T and CD8+T cells using t-test, and FDR is calculated using Benjamini-Hochberg correction. (*: *p*-value < 0.05, **: FDR < 10%). Gene names on the x-axis and asterisks are coloured by significant enrichment in CD4+T cells (blue) or CD8+T cells (orange).

For TCRβ, we observed a strong bias in VDJ gene usage which persisted from the initiation of recombination (DN cells) to the mature T-cell stage (**Fig. 5A and fig. S12**). This bias is not explained by recombination signal sequence (RSS) score (**fig. S13**). The bias does correlate well with genomic position (**fig. S13**), and this is consistent with a looping structure of the locus, which has been observed in mouse (**Fig. 5C**) (*49*). (However, the V gene usage bias that we observe in human is not found in mouse (*25*).) We also observed a preferential association of D2 genes with J2 genes, while D1 genes can recombine with J1 and J2 genes with similar frequency (**fig. S14**). There was no clear association between TCRβ V-D or V-J pairs (**fig. S14A**).

While the initial recombination pattern largely shapes the repertoire, selection also contributes to the preference in TCRβ repertoire. We observed that several TRBV genes were depleted (TRBV6-4, 7-3, 23-1 and 21-1) or enriched (TRBV5-1) after beta-selection compared (DP cells) to before beta-selection (DN cells). Excitingly, this suggests that there are germline-encoded differences between the different Vβ gene’s ability to respond to pMHC stimulation (**fig. S15A**). This result is in line with the molecular finding that Vβ makes the most contacts with pMHC molecule *versus* DJ (and also Vɑ) (*50*).

For the TCRɑ locus, we found a clear association between developmental timing and V-J pairing as described before (*51*): Proximal pairs were recombined first, followed by recombination of distal pairs (**Fig. 5B**), which in turn restricts the pairing between V and J genes (**fig. S14**). This provides direct evidence for progressive recombination of the TCRɑ locus (**Fig. 5D**). Notably, proximal pairs were relatively depleted in mature T cells compared to DP cells, showing a further bias in the positive selection step (**fig. S14B**).

To investigate whether differential TCR repertoire bias exists between cell types, we compared the TCR repertoire of different cell types by running principal component analysis (**Fig. 5E**). Notably, we observed a clear separation of CD8^+^ T cells and other cell types, which was observed across TRBV, TRAV and TRAJ loci. The trend was consistent in all individual donor samples, suggesting that in general, the naive CD8^+^T cell TCR repertoire is skewed by thymic selection. Statistical testing of the difference in odds ratios identified several TCR genes responsible for this phenomenon (**fig. S15B**). The observed trend was largely similar to that seen in naive CD4^+^/CD8^+^T cells isolated from peripheral blood (*22, 23*), providing proof that thymic selection is principally responsible for this bias. Notably, the TRAV-TRAJ repertoire of CD8^+^T cell was biased towards distal V-J pairs compared to other cell types (**Fig. 5F**). Considering that distal repertoires are generated at a later stage of progressive TCRɑ recombination, this suggests slower or less efficient commitment towards the CD8^+^T lineage (**Fig. 5D**). This agrees with finding that CD8 co-receptors are less frequently coupled with Lck kinases in thymus (*52*), providing evidence for thymic regulation of CD8^+^T cell commitment.

Of note, there was also a slight bias towards proximal V-J pairs for CD8ɑɑ^+^T(I) cells that was much more evident in the postnatal thymic sample compared to fetal samples (**fig. S15C**) (*38*). This is in keeping with the inferred trajectory, which showed earlier separation of these cell populations, providing additional support for utilising TRAV-TRAJ proximal-distal bias as a readout for commitment kinetics.

## DISCUSSION

Here we generated a single-cell atlas of the human thymus throughout development *in utero* and postnatal life. We reconstruct the trajectory for human T-cell differentiation combined with TCR repertoire information, which provides valuable molecular insights into T-cell development. Specifically, we dissect multiple steps that result in a bias in the TCR repertoire of mature T cells which persists throughout adult life. As the bias in the TCR repertoire predisposes our reactivity to diverse pMHC combinations, this may have profound implications for how we respond to antigenic challenges. Investigating the biological implications of this bias will shed light on molecular understanding of TCR-pMHC interactions.

Another striking finding is the heterogeneity of unconventional mature T-cell populations in the thymus. We identify GNG4^+^CD8ɑɑ^+^T(I) cells, which constitute a novel CD8ɑɑ^+^ T-cell population specific to the thymus. Previous studies have identified intra-thymic CD8ɑɑ^+^T cells as an intermediate cell state in the thymic cortex that egress from the thymus to populate the intestine as intestinal epithelial lymphocytes (*53*). However, we show that GNG4^+^CD8ɑɑ^+^T(I) cells are located in the thymic medulla and comprise the majority of thymic cells expressing *XCL1*, a chemokine known to recruit XCR1^+^ DCs to the medulla. Therefore, we hypothesize that this is a novel thymus-resident population contributing to thymic function. This is supported by our imaging data showing an interplay of GNG4^+^CD8ɑɑ^+^T(I) cells and DCs in conventional T-cell selection.

The versatile application of T cells as therapeutics in cancer and autoimmune disease is calling for better *in vitro* model systems for T-cell engineering. Our analysis of the thymic microenvironment revealed the complexity of cell types constituting the thymus, and the breadth of interactions between stromal cells and innate immune cells to coordinate thymic development. The intercellular communication network that we describe between thymocytes and supporting cells can form the basis for engineering enhanced *in vitro* culture systems to generate T cells, and inform future T-cell engineering strategies and cell-based clinical therapies.

## Materials and Methods

### Tissue Acquisition

All tissue samples used for this study were obtained with written informed consent from all participants in accordance with the guidelines in The Declaration of Helsinki 2000 from multiple centres. Human fetal tissues were obtained from the MRC/Wellcome Trust-funded Human Developmental Biology Resource (HDBR, http://www.hdbr.org) with appropriate maternal written consent and approval from the Newcastle and North Tyneside NHS Health Authority Joint Ethics Committee (08/H0906/21+5). HDBR is regulated by the UK Human Tissue Authority (HTA; www.hta.gov.uk) and operates in accordance with the relevant HTA Codes of Practice. Human paediatric samples were obtained from Ghent University Hospital with appropriate written consent and approval from the Ghent University Hospital Ethics Committee (B670201319452). The human adult deceased donor sample was obtained from the Cambridge Biorepository for Translational Medicine (CBTM) with appropriate written consent and approval from the Cambridge University Ethics Committee (reference 15/EE/0152, East of England Cambridge South Research Ethics Committee).

### Tissue Processing

All tissues were processed immediately after isolation using a consistent protocol with slight modification. Tissue was transferred to a sterile 10mm^2^ tissue culture dish and cut into <1mm^3^ segments before being transferred to a 50mL conical tube. Fetal and paediatric tissues were digested with 1.6mg/mL collagenase type IV (Worthington) in RPMI (Sigma-Aldrich) supplemented with 10%(v/v) heat-inactivated fetal bovine serum (Gibco), 100U/mL penicillin (Sigma-Aldrich), 0.1mg/mL streptomycin (Sigma-Aldrich), and 2mM L-Glutamine (Sigma-Aldrich) for 30 minutes at 37°C with intermittent shaking. Adult organ donor sample was digested with 0.2 mg/ml Liberase ™ (Roche)/0.125 KU DNase1 (Sigma-Aldrich)/10mM HEPES in RPMI for 30 minutes at 37°C with intermittent shaking. Digested tissue was passed through a 100µm filter, and cells collected by centrifugation (500g for 5 minutes at 4°C). For fetal and adult organ donor samples, cells were treated with 1X red blood cell (RBC lysis buffer (eBioscience) for 5 minutes at room temperature and washed once with flow buffer (PBS containing 5%(v/v) FBS and 2mM EDTA) prior to cell counting. For paediatric samples, RBC lysis step was omitted.

### Fetal developmental stage assignment and chromosomal assessment

Embryos up to 8 post conception weeks (PCW) were staged using the Carnegie staging method (*54*). After 8 PCW, developmental age was estimated from measurements of foot length and heel to knee length and compared against a standard growth chart (*55*). A piece of skin, or where this was not possible, chorionic villi tissue, was collected from every sample for Quantitative Fluorescence-Polymerase Chain Reaction analysis using markers for the sex chromosomes and the following autosomes: 13, 15, 16, 18, 21, 22, which are the most commonly seen chromosomal abnormalities.

### Flow cytometry and FACS for Single-cell RNA Sequencing

Isolated thymus cells were stained with a panel of antibodies prior to sorting based on CD45 or CD3 expression gate. The anti-human monoclonal antibodies used for flow cytometry based immunophenotyping and sorting are listed in Table S1. An antibody cocktail was freshly prepared by adding 3µL of each antibody in 50µL Brilliant Stain Buffer (BD) per tissue. Cells (≤10×10^6^) were resuspended in 50-100µL flow buffer and an equal volume of antibody mix was added to cells from each tissue. Cells were stained for 30 minutes on ice, washed with flow buffer and resuspended at 10×10^6^ cells/mL. DAPI (Sigma-Aldrich) was added to a final concentration of 3µM immediately prior to sorting. Flow sorting was performed on a BD FACSAria™ Fusion instrument using DIVA V8, and data analysed using FlowJo V10.4.1. Cells were gated to remove dead cells and doublets, and then sorted for 10X or SS2 scRNAseq analysis. For 10X droplet microfluidic analysis, cells were sorted into chilled FACS tubes coated with FBS and prefilled with 500µL sterile PBS. Paediatric samples were sorted into 50% FCS and 50% IMDM medium (supplemented with 1% glu, 1%P/S and 10% FCS). For SS2 scRNAseq analysis, single cells were index-sorted into 96-well lo-bind plates (Eppendorf) containing 10µL lysis buffer (TCL 858 (Qiagen) + 1% (v/v) E-mercaptoethanol) per well.

### Single molecule RNA FISH

Samples were fixed in 10% NBF, dehydrated through an ethanol series and embedded in paraffin wax. Five-micrometre samples were cut, baked at 60 °C for 1 h and processed using standard pre-treatment conditions, as per the RNAScope multiplex fluorescent reagent kit version 2 assay protocol (manual) or the RNAScope 2.5 LS fluorescent multiplex assay (automated). The RNAScope probes used for this study are listed in Table S2. TSA-plus fluorescein, Cy3 and Cy5 fluorophores were used at 1:1500 dilution for the manual assay, or 1:300 dilution for the automated assay. Slides were imaged on different microscopes: Hamamatsu Nanozoomer S60 or 3DHistech Pannoramic MIDI. Filter details were as follows: DAPI: excitation 370–400, BS 394, emission 460–500; FITC: excitation 450–488, BS 490, emission 500–55; Cy3: excitation 540–570, BS 573, emission 540–570; Cy5: excitation 615–648, BS 691, emission 662–756.

### Thymic fibroblasts culture derivation and phenotypic characterisation

Thymic explants were derived from foetal biopsies at different thymic stages (HDBR Newcastle University - NewCcstle Upon Tyne, REC reference: 19/NE/0290 and HDBR University College of London - London, REC reference: 18/LO/0822) and cultured on a precoated Matrigel (Corning) 6mm dish in DMEM (Life Technologies) supplemented with 15% FBS HI (Life Tehnologies) + 1% Pen/Strept (Sigma-Aldrich), 1% L-glutamine (Life Technologies), 1% Non-Essential Aminoacids (Life Technologies) and 100mM beta-Mercaptoethanol (Life Technologies). Fibroblast cells come out of explants at around 7 days of culture and are left on the plate until outgrowths are confluent enough to pass. The culture is therefore kept for 5-6 passages and phenotypic analysis was performed at multiple passages.

Fibroblasts were detached with trypsin 1X (Sigma-Aldrich) for 3’ at 37C and subsequently resuspended in completed media before collection. Cells are harvested and phenotypic analysis is performed on 500,000 cells per sample. Cells were stained at 4°C for 30 min in Hanks Balanced Salt Solution-2% FBS with the following markers: anti-THY1 AF700 1:100 (Biolegend), anti-PDGFRalpha PE 1:100 (Biolegend) and PI-16 (BD) 1:50. Cells are washed in an excess of HBSS + 2%FBS and are resuspended in HBSS + 2%FBS with DAPI (Sigma-Aldrich) to discriminate live from dead cells.

### Library Preparation and Sequencing

For the droplet-encapsulation scRNA-seq experiments, 8000 live, single, CD45^+^ or CD45^-^ FACS-isolated cells were loaded on to each of the Chromium Controller (10x Genomics). Single cell cDNA synthesis, amplification and sequencing libraries were generated using the Single Cell 3’ and 5’ Reagent Kit following the manufacturer’s instructions. The libraries from up to eight loaded channels were multiplexed together and sequenced on an Illumina HiSeq 4000. The libraries were distributed over eight lanes per flow cell and sequenced using the following parameters: Read1: 26 cycles, i7: 8 cycles, i5: 0 cycles; Read2: 98 cycles to generate 75bp paired end reads.

For the plate-based scRNA-seq experiments, a slightly modified Smart-Seq2 protocol was used as previously described (*42*). After cDNA generation, libraries were prepared (384 cells per library) using the Illumina Nextera XT kit. Index v2 sets A, B, C and D were used per library to barcode each cell for multiplexing. Each library was sequenced (384 cells) per lane at a sequencing depth of 1-2 million reads per cell on HiSeq 4000 using v4 SBS chemistry to create 75bp paired end reads.

### Alignment, quantification and quality control of single cell RNA sequencing data

Droplet-based sequencing data was aligned and quantified using the Cell Ranger Single-Cell Software Suite (version 2.0.2 for 3’ chemistry and version 2.1.0 for 5’ chemistry, 10x Genomics Inc) using the GRCh38 human reference genome (official Cell Ranger reference, version 1.2.0). Cells with fewer than 2000 UMI counts and 500 detected genes were considered as empty droplets and removed from the dataset. Cells with more than 7000 detected genes were considered as potential doublets and and removed from the dataset.

Smart-seq2 sequencing data was aligned with *STAR* (version 2.5.1b), using the STAR index and annotation from the same reference as the 10x data. Gene-specific read counts were calculated using *htseq-count* (version 0.10.0). Scanpy (version 1.3.4) python package was used to load the cell-gene count matrix and perform downstream analysis.

### Doublet detection

To exclude doublets from single-cell RNA sequencing data, we applied scrublet (*56*) algorithm per sample to calculate scrublet-predicted doublet score per cell with following parameters: sim_doublet_ratio = 2; n_neighbors=30; expected_doublet_rate= 0.1. Any cell with scrublet score > 0.7 was flagged as doublet. To propagate the doublet detection into potential false-negatives from scrublet analysis, we over-clustered the dataset (*sc.tl.louvain* function from scanpy package version 1.3.4; resolution = 20), and calculated the average doublet score within each cluster. Any cluster with averaged scrublet score > 0.6 was flagged as a doublet cluster. All remaining cell clusters were further examined to detect potential false-negatives from scrublet analysis according to the following criteria: (1) Expression of marker genes from two distinct cell types which are unlikely according to prior knowledge (i.e. *CD3* for T cells and *CD19* for B cells), (2) Higher number of UMI counts and (3) Lack of unique marker gene defining the cluster. Finally, all flagged doublets are clustered within themselves, and these doublet clusters are used to train logistic regression model (*sklearn.linear_model.LogisticRegression*; penalty = ‘l2’, C=0.2) together with annotated cell types, and the doublets predicted by this model are also flagged as doublets. All clusters flagged as doublets were removed from further downstream biological analysis.

### Clustering and annotation of scRNA-seq data

Downstream analysis included data normalisation (*scanpy.api.pp.normalize_per_cell* method, scaling factor 10000), log-transformation (*scanpy.api.pp.log1p),* variable gene detection (*scanpy.api.pp.filter_gene_dispersion*), data feature scaling (*scanpy.api.pp.scale*), PCA analysis (*scanpy.api.pp.pca*, from variable genes), batch-balanced neighbourhood graph building (*scanpy.api.pp.bbknn*) and Louvain graph-based clustering (*scanpy.api.tl.louvain*, clustering resolution manually tuned) performed using the python package scanpy (version 1.3.4). Custom defined cell cycle gene sets (**Table S3**) were removed from the list of variable genes to remove cell-cycle associated variation. Cluster cell identity was assigned by manual annotation using known marker genes as well as computed differentially expressed genes (DEGs) using custom python function. Clusters with clear and uniform identity were annotated first, and a logistic regression model was trained based on this annotation. This model was used to predict the identity of cells in a cluster with a mixture of different cell types, which can be computationally clustered together due to transcriptional similarity. To achieve a high resolution annotation, we separated broadly annotated cells (e.g. Epithelial cells, single positive T cells) and repeated the procedure of variable gene selection, which allowed the annotation of smaller and fine-grained cell subsets (e.g. mTECs, regulatory T cells).

### Alignment of data across different batches

Batches for batch alignment can come from different chemistries used on the same set of cells, e.g. 10X chemistry (5’ and 3’), or from cells from different donors analysed using the same chemistry. In other words, there can be technical or biological differences between batches. We performed iterative batch correction, first by roughly aligning batches across similar samples (e.g. all foetal samples or paediatric samples) using *scanpy.api.pp.bbknn* function. We used this batch-aligned manifold to annotate cell types. After achieving a coarse-grained cell type annotation, we fitted a linear model using batches (e.g. 10X chemistry, donors) or cell type annotation as a categorical variable. Then we regressed out variations explained by batch variables, and kept residuals, which contain biological information. After this, we aligned batches again using the *scanpy.api.pp.bbknn* function to achieve a high-resolution and batch-mixed manifold, which is used for refining annotation, visualisation and trajectory analysis.

### Estimating cellular composition per sample

To estimate the relative proportion of each cell type in different samples, we defined broad categories of cell types (e.g. lymphocytes, myeloid cells, total cells), and calculated the proportions of each cell type within selected group of cells. If all cell types used for a comparison come from the same sorting gate, we simply calculated the proportion as: number of cells in specific cell type / total number of cells in comparison set. When cell types used for comparison are derived from multiple sorting gates, we calculated a normalisation factor for each sorting gate as: number of cells sorted in a specific sorting gate / total number of sorted cells across multiple sorting gates, and multiplied this normalisation factor to the number of cells in each sorting gate. These normalised numbers are used to calculate proportions, which eliminates bias caused by sorting different number of cells into different gates.

### Trajectory analysis

To model differentiation trajectories, a combination of linear regression and batch-alignment algorithms were applied as described above to generate a neighbourhood graph. The robustness and accuracy of batch-alignment was tested by comparing multiple batch-alignment methods. Among the resulting manifolds, we selected the one with the best fit to well-known sequential events in T-cell differentiation such as TCR recombination. We then calculated diffusion pseudotime (*57*) using the *scanpy.api.tl.dpt* function in scanpy, which starts from the manually selected progenitor cell. The progenitor cell is selected from the extremities of diffusion components. Cells are binned based on the pseudotime ordering, and differentially expressed genes are identified as genes whose expression is significantly different from the randomly permuted background in any of the bins.

### Visualisation of the transcription factor network

Transcription factor network analysis was performed as previously described (*26*). First, gene expression levels were imputed by taking an average of 30-nearest neighbors in three-dimensional UMAP space. An annotation score for each cell type was calculated by measuring the frequency of cell types amongst the 30-nearest neighbors which are used for imputation. To remove redundant information, cells were randomly sampled from each unit voxel from the three-dimensional UMAP space. The human transcription factors were selected from AnimalTFDB3 (*58*). Only highly-variable transcription factors were subject to calculation of the correlation matrix, which was subsequently used for graph building and visualisation using the force-directed graph function implemented in the scanpy package.

### TCR VDJ sequence analysis

10X TCR-enriched libraries are mapped with the Cell Ranger Single-Cell Software Suite (version 2.1.0, 10x Genomics Inc) to the custom reference provided by the manufacturer (version 2.0.0 GRCh38 VDJ reference). VDJ sequence information was extracted from the output file “filtered_contig_annotations.csv.” The merged VDJ output dataset is available in our data repository (see Data and materials availability). Chains which contained full-length recombinant sequence and supported by more than 2 UMI counts were selected, and linked to the cellular transcriptome data based on cell barcodes. These chains were considered as productive if a functional ORF covering the CDR3 region could be found. To compare V, D, J gene usage per cell type, each V, D, J gene count in each specific cell type was normalised by the sum of counts within that cell type, and then converted to a z-score per gene. Student’s t-test was used to compare the z-scores between different cell types. Cochran–Mantel–Haenszel test was also used to compare profiles between CD4+T and CD8+T cells, which yielded comparable results.

### Cell-cell interaction analysis

Specific interactions between cells are modeled using CellPhoneDB (www.CellPhoneDB.org) as previously described (*35*). To minimise computational burden and equally represent different cell types, we downsampled the dataset by randomly sampling 1000 cells from each cell type. We modified cell-cell interaction scores by multiplying average expression level of each ligand and receptor gene within cell-cell pairs, and maximum-normalising this score. The list of chemokines was retrieved from the HUGO Gene Nomenclature Committee.

## Acknowledgements

We gratefully acknowledge the Sanger Flow Cytometry Facility, Newcastle University Flow Cytometry Core Facility, Sanger Cellular Generation and Phenotyping (CGaP) Core Facility, and Sanger Core Sequencing pipeline for support with sample processing and sequencing library preparation. We thank the MRC/Engineering and Physical Sciences Research Council Newcastle Molecular Pathology Node for support on paraffin embedding fetal tissues. We thank Jana Eliasova for graphical images and Jinwook Choi for helpful discussions, and Sarah Aldridge for editing the manuscript. The human embryonic and fetal material was provided by the Joint MRC / Wellcome (MR/R006237/1) Human Developmental Biology Resource (www.hdbr.org). The material from the deceased organ donor was provided by the Cambridge Biorepository for Translational Medicine. Paediatric thymus material was provided by Ghent University Hospital. We are grateful to the donors and donor families for granting access to the tissue samples.

## Author contributions

J-E.P, S.A.T, M.H, and T.T. designed the experiments. J-E.P; R.A.B; D.M.P.; E.S.; M.L.; C.D.C; J.R.F; N.V.; K.M.; performed sampling and library prep with help from S.W.; D.M.; A.F.; A.F.; L.M.; G.R.; D.D.; R.V-T.. J-E.P.; D.J.K; K.P.; M.L; C.D.C analysed the data. J-E.P.; C.D.C; E.T.; A.W-C.; R.A.B; R.R. performed validation experiments. J-E.P; C.D.C; S.A.T; M.H.; T.T. wrote the manuscript with contributions from R.A.B; K.B.M; Y.S.; M.R.C.; P.B; S.B.; R.V-T.; D.J.K. All authors read and accepted the manuscript.

## Data and materials availability

All open access-consented sequencing data will be deposited in the Human Cell Atlas Data Coordination Portal upon acceptance of the manuscript (i.e. all embryonic, fetal and organ donor-derived data).

---

**Fig. S1.**
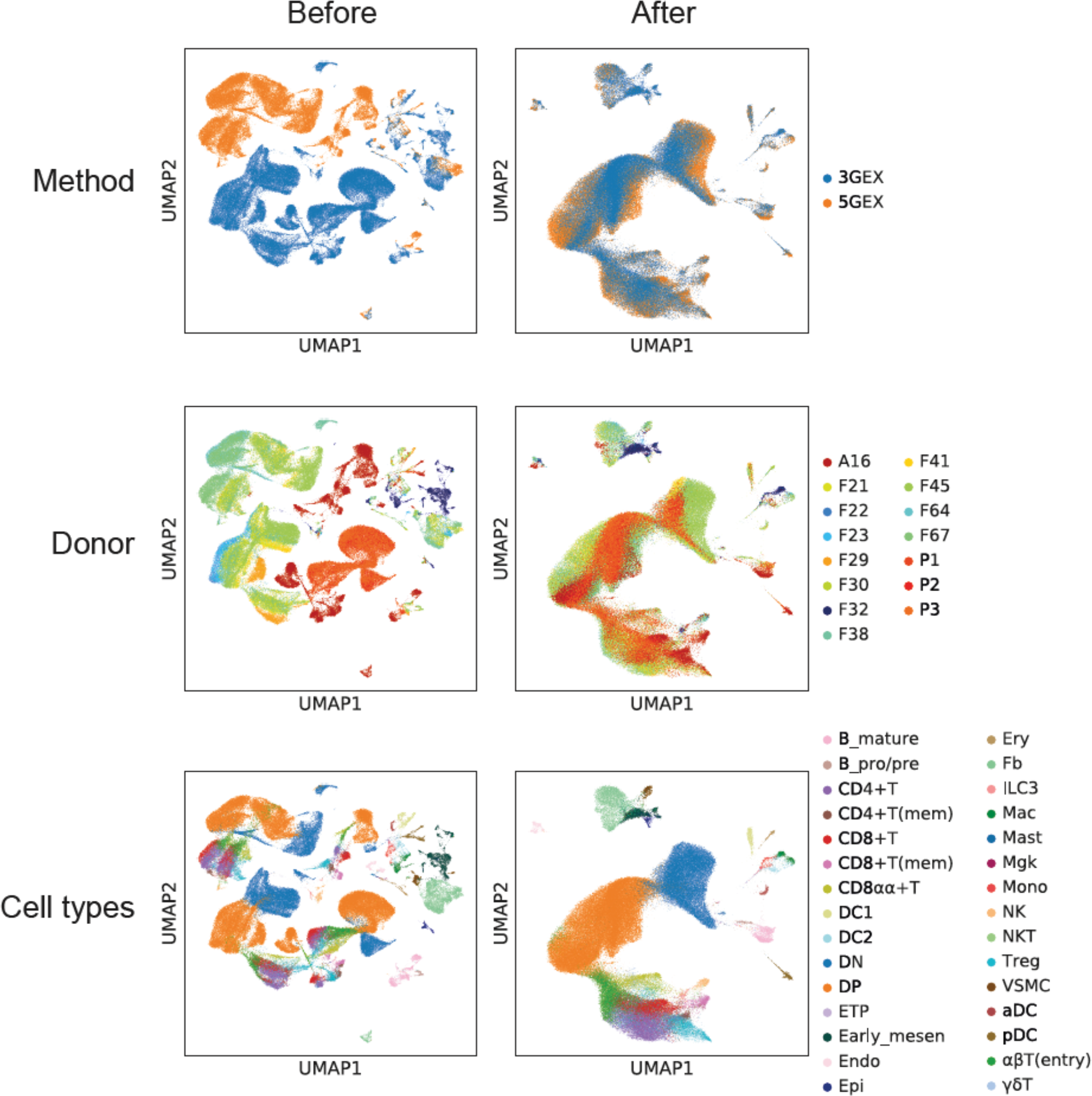
UMAP visualisation of entire dataset before (left) and after (right) batch alignment. Cells are coloured by methods (top), donors (middle) and cell types (bottom).

**Fig. S2.**
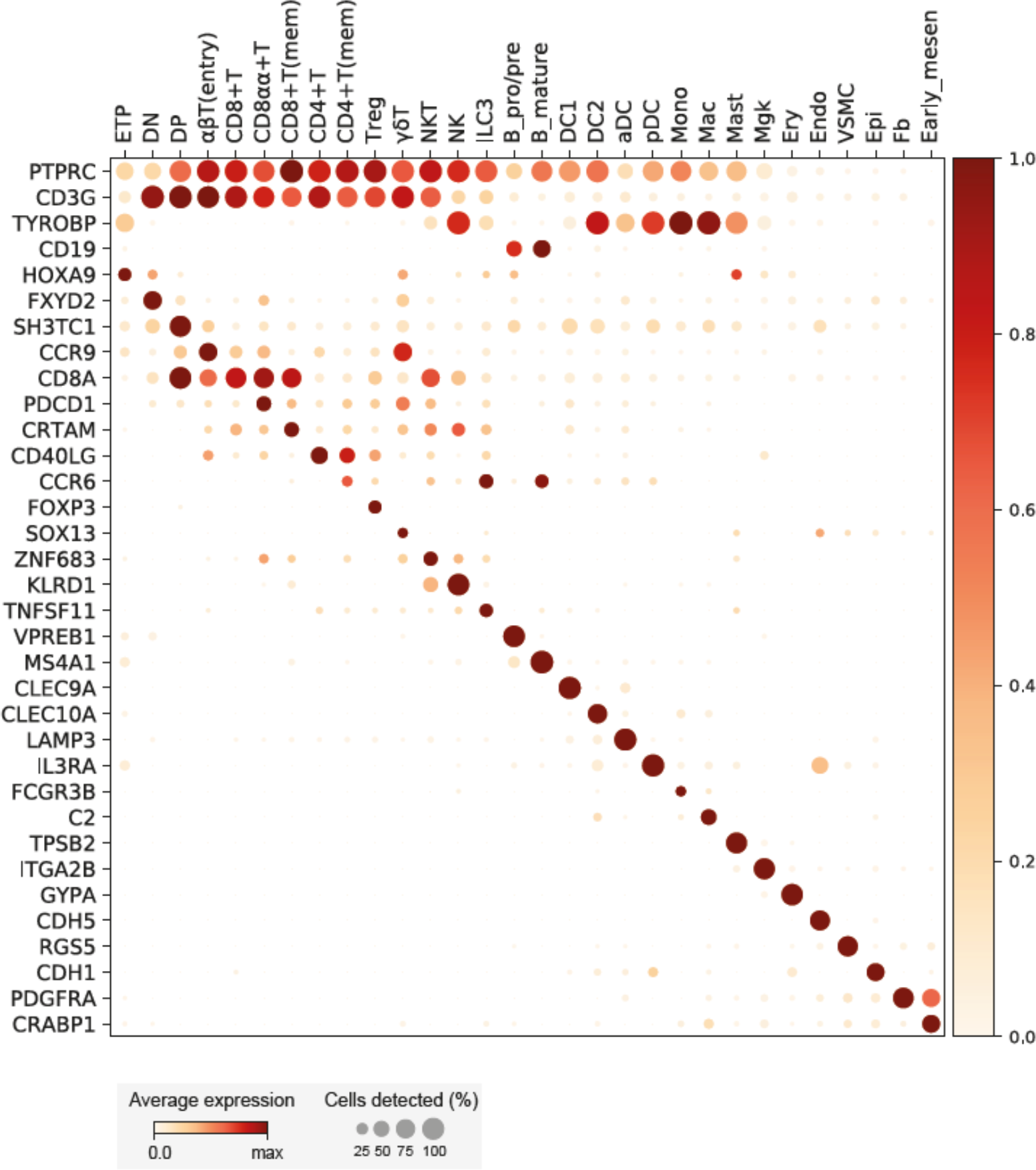
Dot plot showing marker gene expression for annotated cell types. ETP: early thymic progenitors, DN: double negative T cells, DP: double positive T cells, Treg: regulatory T cells, ILC3: innate lymphoid cell type 3, B_pro/pre: pro-B cells and pre-B cells, DC1: conventional dendritic cell type 1, DC2: conventional dendritic cell type 2, aDC: activated dendritic cells, pDC: plasmacytoid dendritic cells, Mono: monocytes, Mac: macrophage, Mast: mast cells, Mgk: megakaryocytes, Ery: erythrocytes, Endo: endothelial cells, VSMC: vesicular smooth muscle cells, Epi: epithelial cells, Fb: fibroblasts, Early_mesen: mesenchymal cells in 7 PCW fetus

**Fig. S3.**
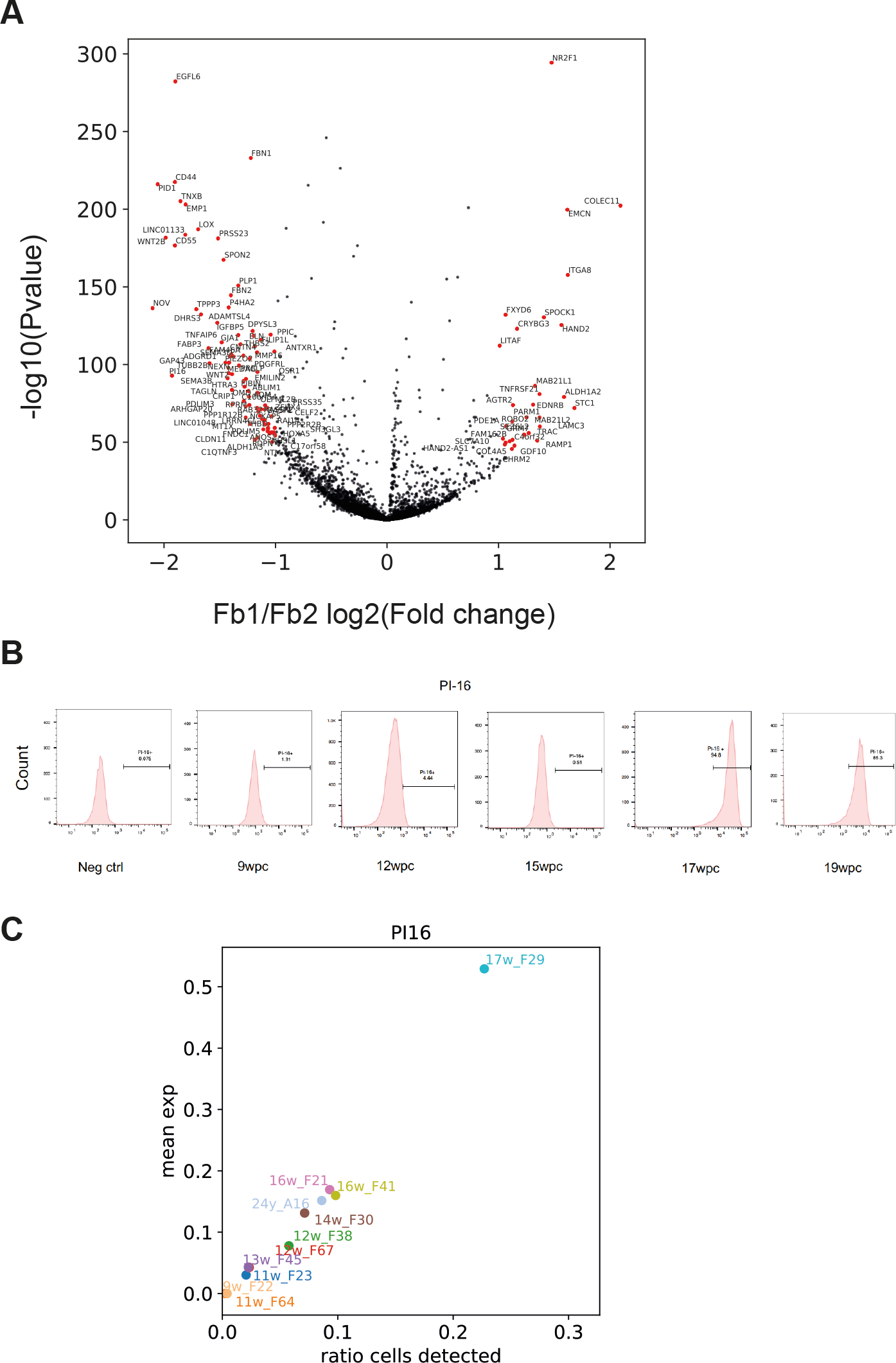
(A) Volcano plot showing differentially expressed genes between two types of thymic fibroblasts. X-axis and y-axis represent log2(fold change) and -log10(p-value) respectively. **(B)** FACS analysis of PI protein level in thymic fibroblast explant culture from different stage of human fetal thymus. **(C)** Expression level of PI mRNA level in single-cell RNA sequencing data from different stage of human fetal thymus

**Fig. S4.**
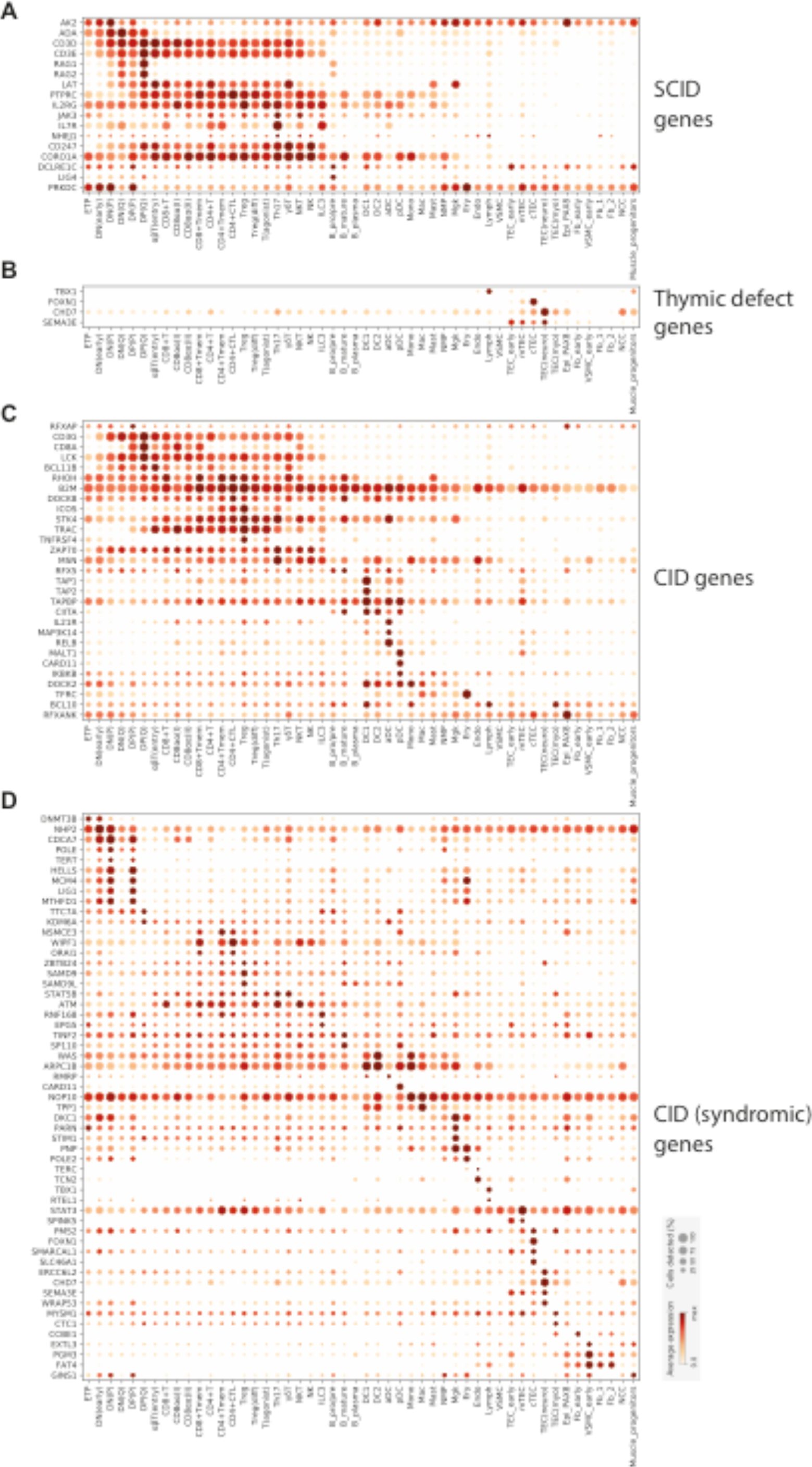
Dot plot showing the expression of genes causing Severe Combined Immunodeficiency (SCID) (A), thymic defects (B), Combined Immunodeficiency (CID) (C), and syndromic CID (D). Genes are taken from the IUIS Classification of Inborn Errors of immunity (February 2018). DN(early): double negative, committed T cells, NMP: neutrophil-myeloid progenitors, Lymph: lymphatic endothelial cells, TEC_early: undifferentiated thymic epithelial cells (TECs) in 7 PCW fetus, mTEC: medullary TECs, cTEC: cortical TECs, Fb_early: undifferentiated fibroblasts in 7 PCW fetus, Fb_1: fibroblast type 1, Fb_2: fibroblast type 2, NCC: neural crest cells in 7 PCW fetus, Muscle_progenitors: muscle progenitors in 7 PCW fetus. Other abbreviations are defined in Fig. S2.

**Fig. S5.**
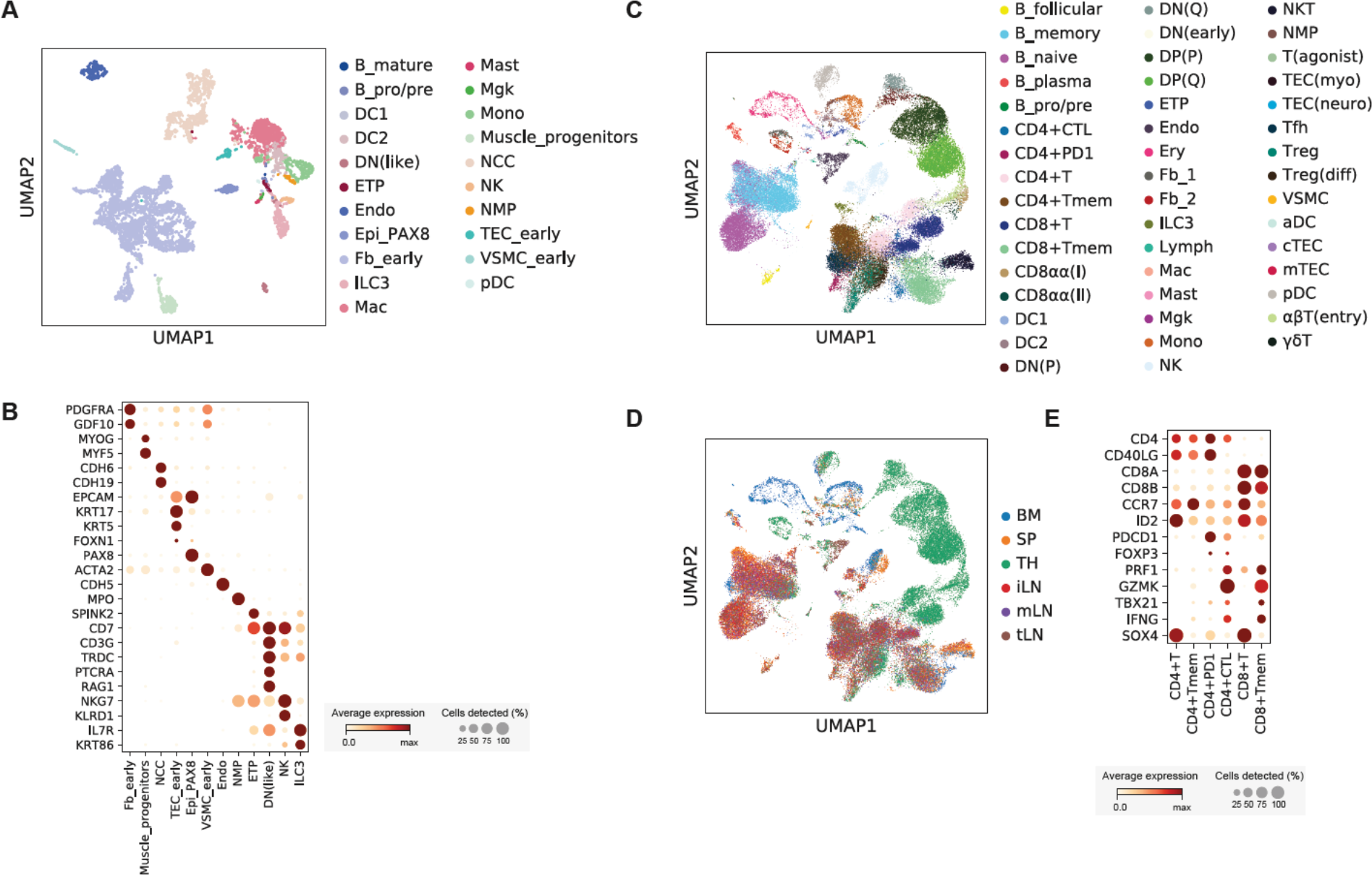
(A) UMAP plot showing cell type annotations for 7 PCW thymus sample. **(B)** Dot plot showing marker gene expression for 7 PCW thymus sample. **(C)** UMAP plot showing cell type annotations for the young adult sample (20-25 years old) which have morphological signature of degeneration. **(D)** Organ composition for UMAP plot shown in (C). **(E)** Dot plot showing marker gene expression for mature T cells found in young adult sample. Abbreviations are as defined from Fig. S2 and S3.

**Fig. S6.**
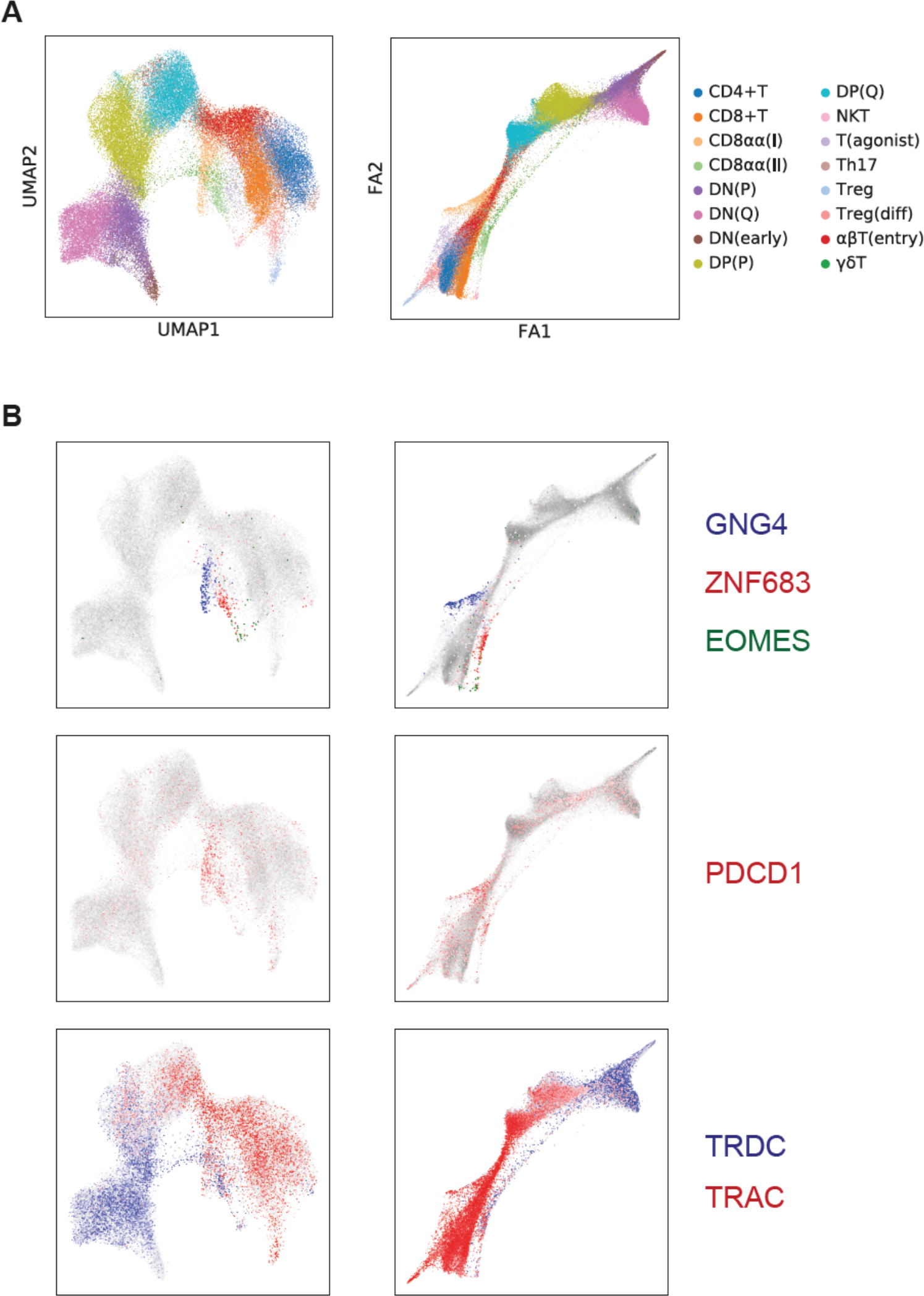
(A) UMAP plot (left) and force directed graph plot (right) showing T cell development trajectory. **(B)** UMAP plot (left) and force directed graph plot (right) showing marker gene expression for CD8ɑɑ^+^ T subtypes in thymus.

**Fig. S7.**
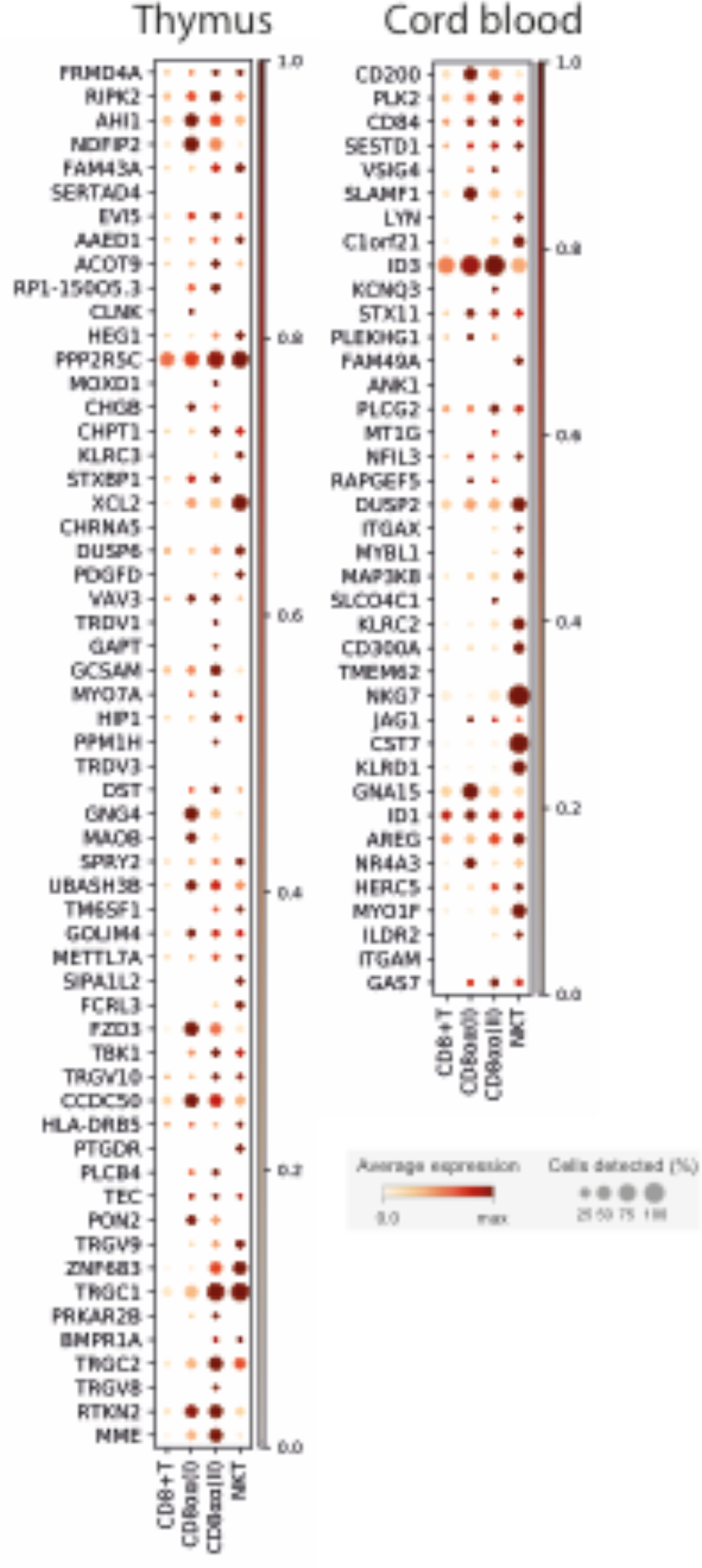
Dot plot showing expression level of CD8ɑɑ^+^ T marker genes enriched in thymus (left) or cord blood (right) across conventional CD8^+^ T cells and three CD8ɑɑ^+^ T types found in thymus.

**Fig. S8.**
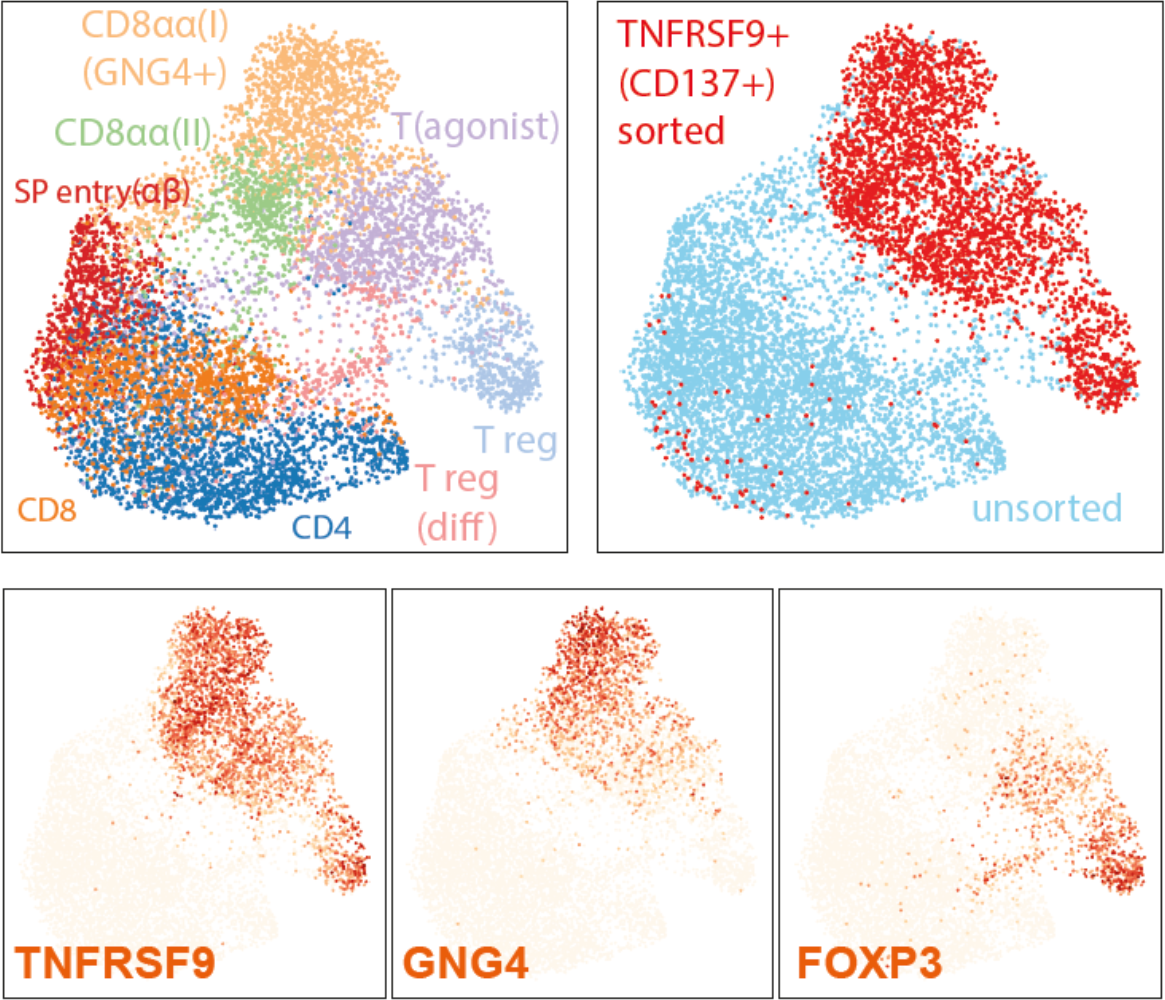
UMAP plot showing CD137+CD3+ sorted population from 12 PCW fetal thymus. Sorted cells (top right, red) were compared to unsorted mature T cells (top right, skyblue) from the same individual. Gene expression of CD8ɑɑ+T(I) marker (*GNG4*), Treg marker (*FOXP3*) and marker shared between these two groups (*TNFRSF9/CD137*) are shown.

**Fig. S9.**
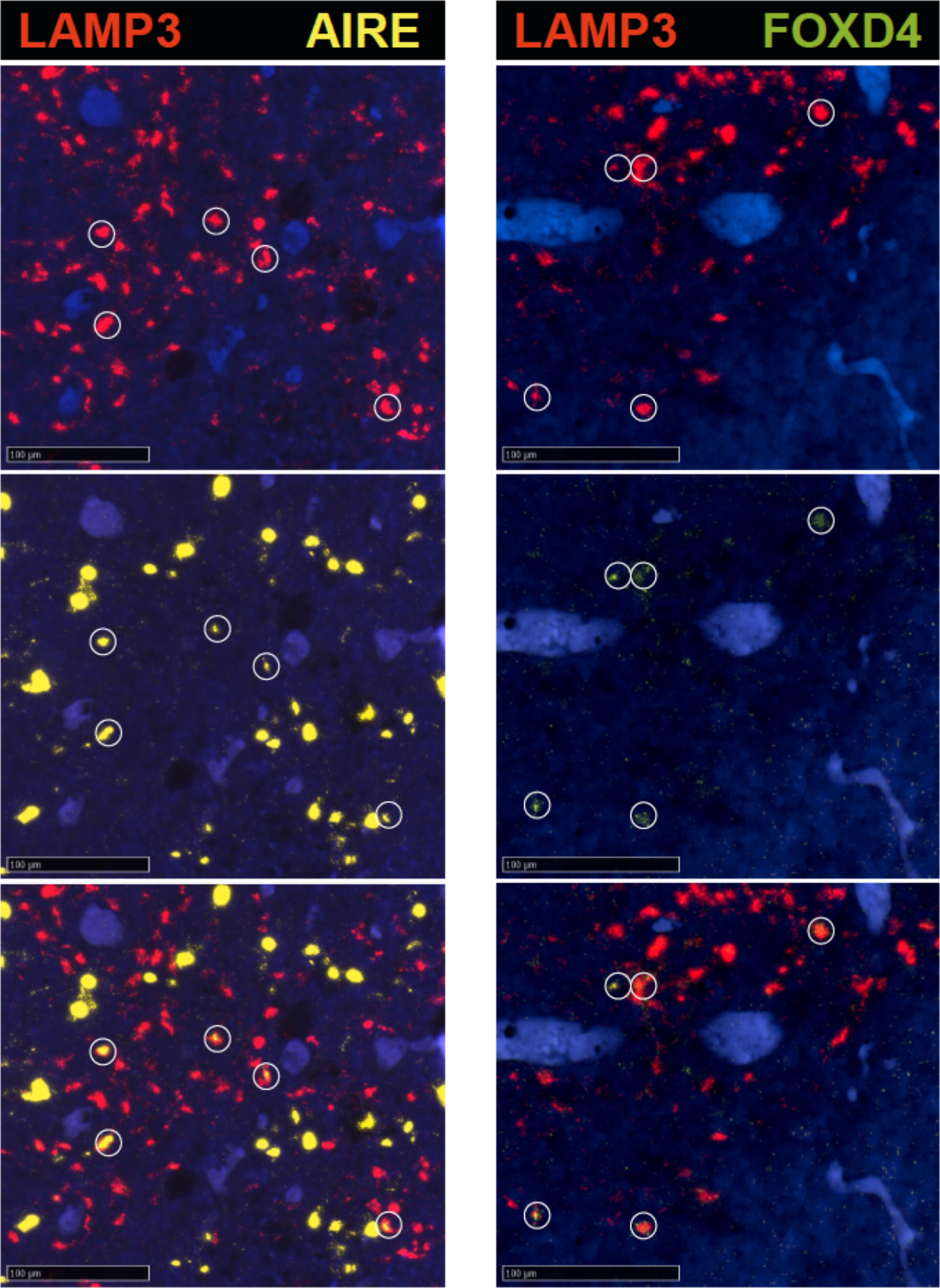
RNA single-molecule FISH detection of various genes expressed in aDCs (*LAMP3*, *AIRE*, *FOXD4*) on 15 PCW fetal thymus tissue section. Cells with expression of both genes are marked with circle.

**Fig. S10.**
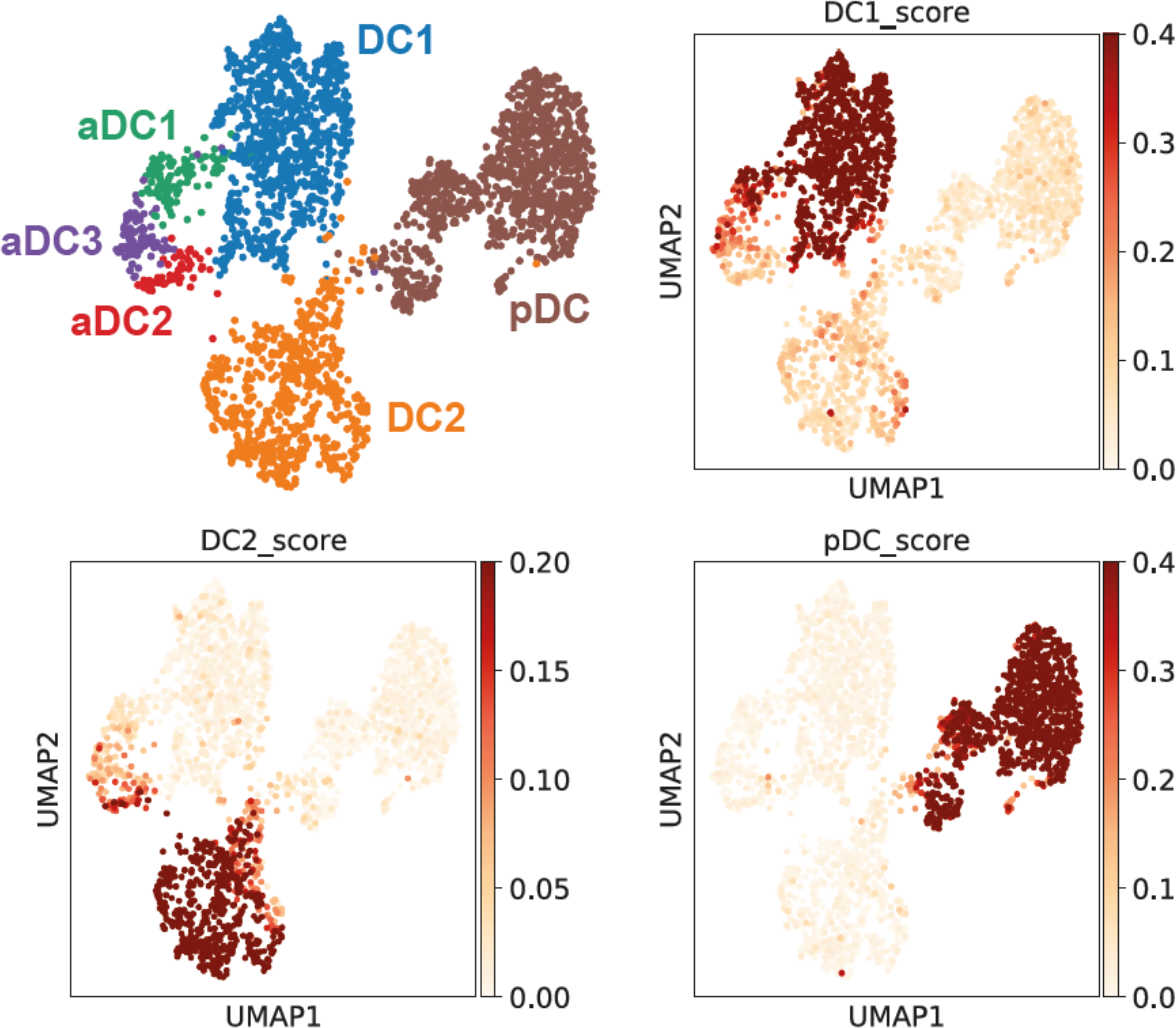
UMAP plot showing DC subtypes found in human thymus (top left). The same UMAP plot is used to show the cells with high DC1, DC2 and pDC scores, which are calculated by taking average of expression level for lineage specific genes.

**Fig. S11.**
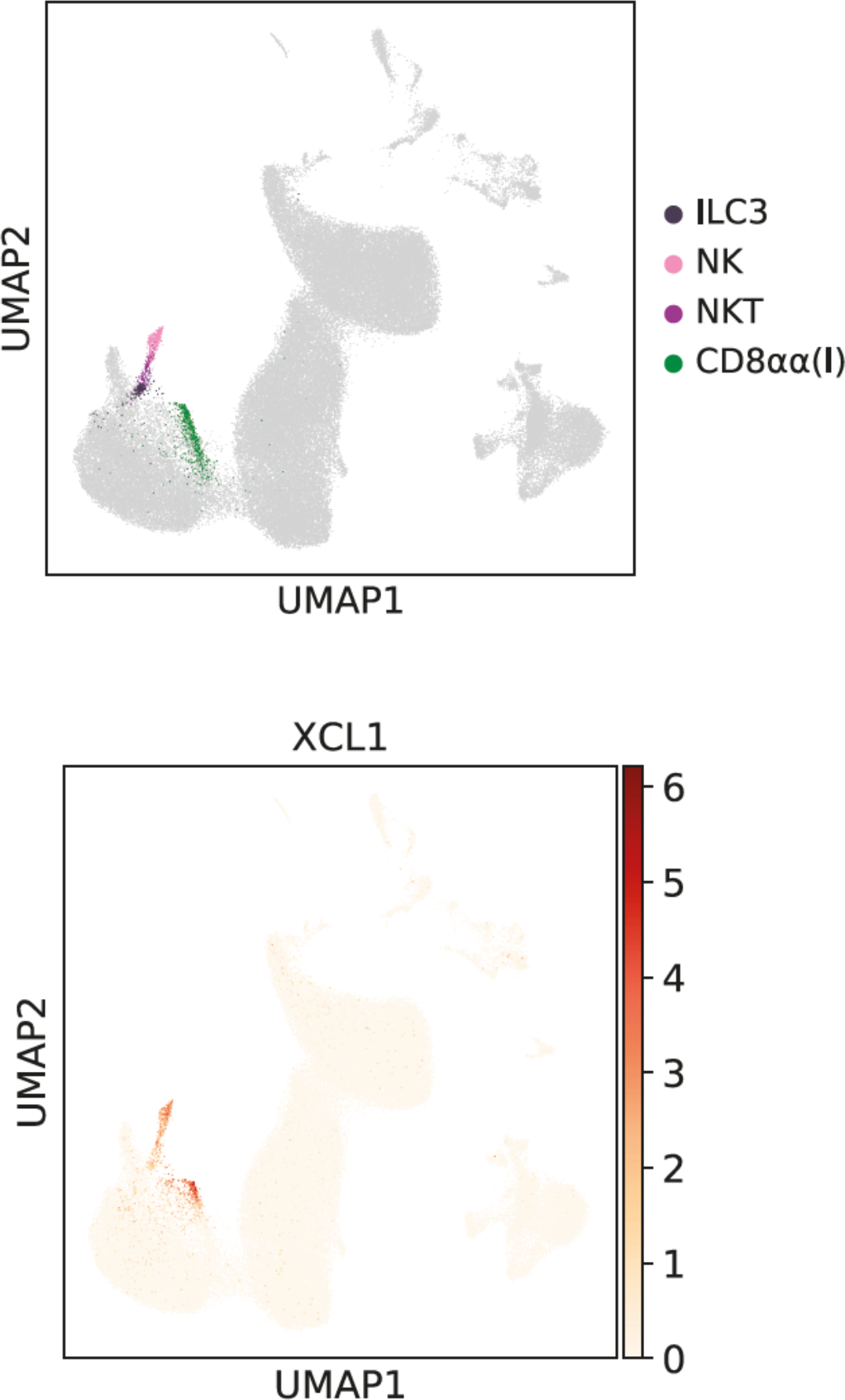
UMAP plot showing XCL1 expression level in fetal thymus (top) and cell types expressing XCL1 (bottom)

**Fig. S12.**
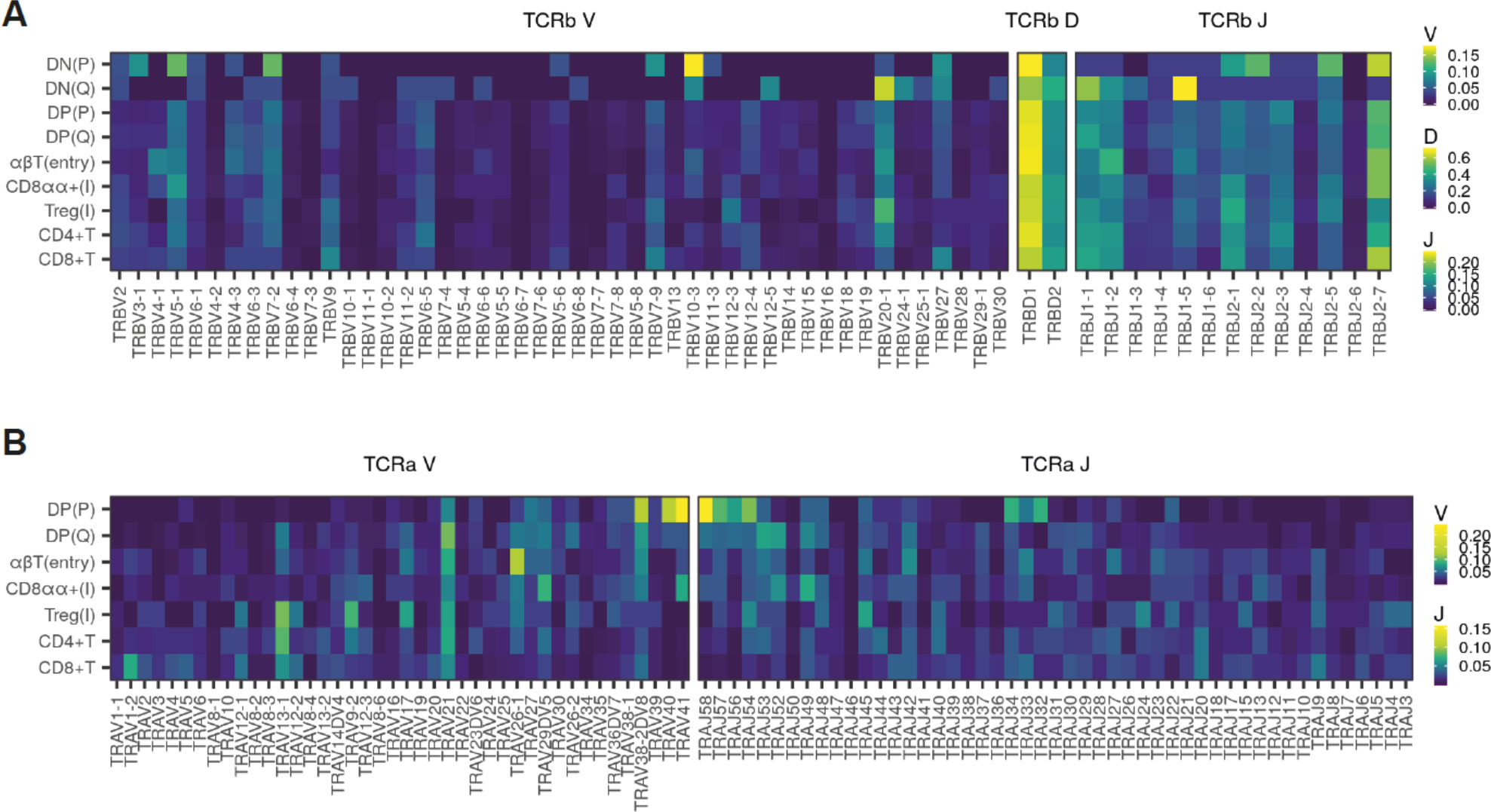
(A) Heatmap showing the proportion of TCRβ V, D, J gene segment present at different stages of T cell development for the young adult sample (20-25 years old). Gene segments are positioned according to genomic location. **(B)** Same scheme applied to TCRα V and J gene segments.

**Fig. S13.**
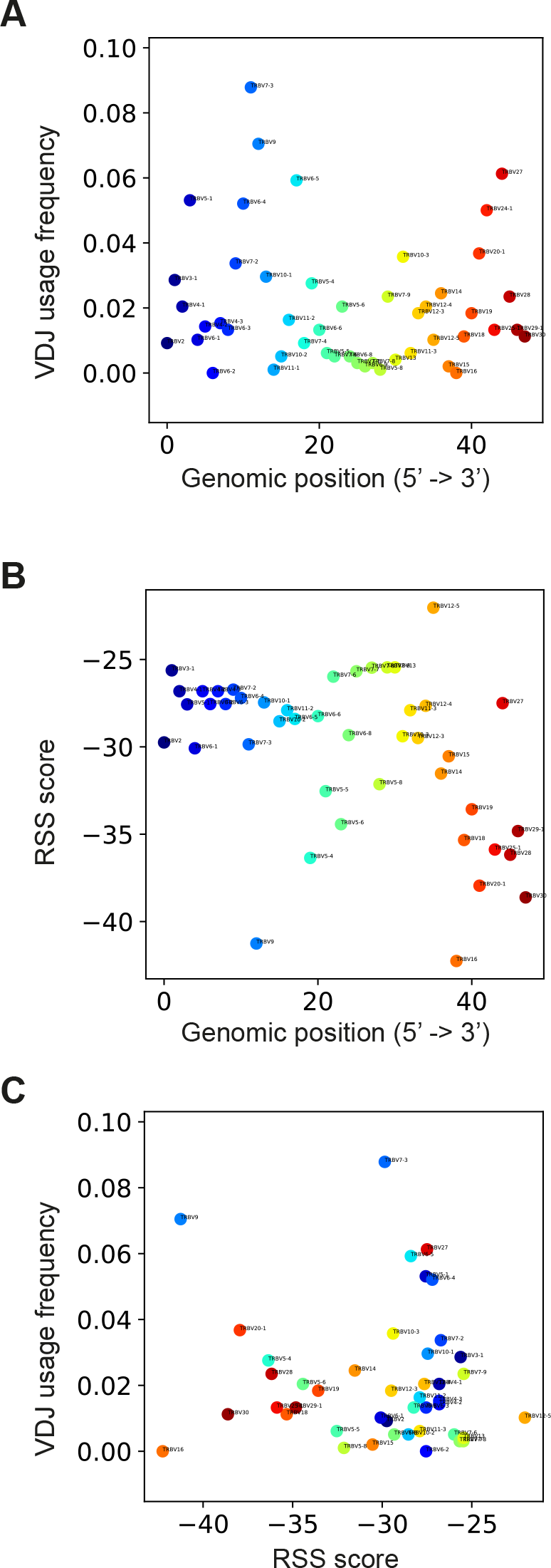
(A) Scatter plot comparing genomic position (x-axis) and relative usage (y-axis) for TCRβ V genes. Genes are coloured based on genomic position. The same colour scheme is applied for following figures. **(B)** Scatter plot comparing genomic position (x-axis) and RSS score (y-axis) for TCRβ V genes. **(C)** Scatter plot comparing RSS score (x-axis) and relative usage (y-axis) for TCRβ V genes.

**Fig. S14.**
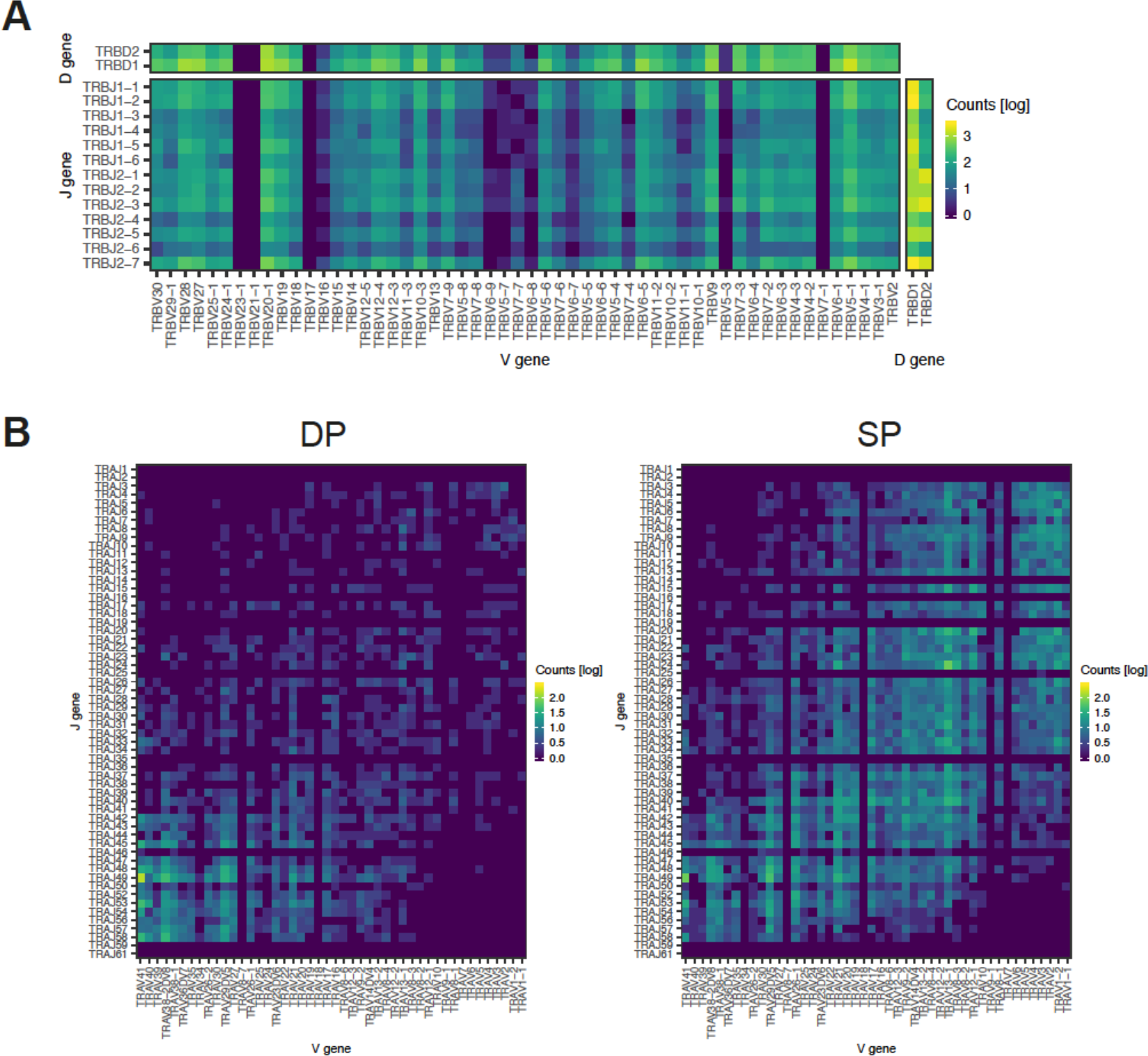
(A) Relative frequency (log scale) of V-J, V-D, J-D gene pairs in TCRβ locus. **(B)** Relative frequency (log scale) of V-J gene pairs in TCRɑ locus. Dataset is divided into DP and SP stages to highlight the enrichment of proximal pairs in DP stage.

**Fig. S15.**
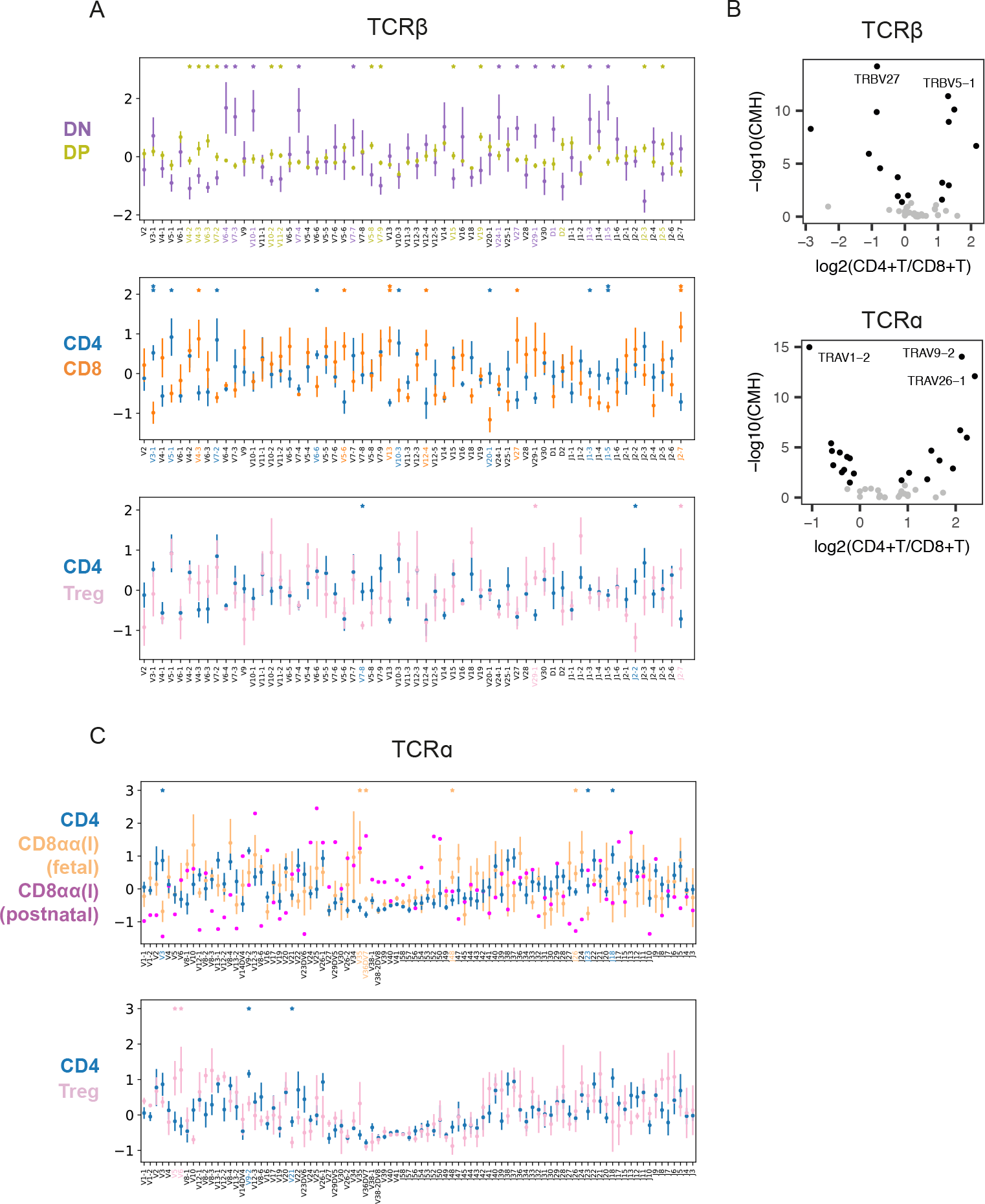
(A, B) Relative usage of V, D and J gene segments according to cell types for TCRβ locus (A) or TCRɑ locus (B). *Z*-score for each segment is calculated from the distribution of normalised proportions stratified by the cell type and sample. *P*-value is calculated by comparing z-scores using t-test, and FDR is calculated using Benjamini-Hochberg correction. (*: *p*-value < 0.05, **: FDR < 10%) Gene names on the x-axis and asterisks are coloured by significant enrichment. For CD4 vs CD8ɑɑ+T (I) comparison, CD8ɑɑ+T (I) data points are separated into fetal samples (n=4) and post-natal sample (n=1, young adult) to highlight differences between fetal sampels and young adult sample. All other comparisons are inclusive of both fetal and post-natal samples. Consistency between fetal and post-natal samples are separately confirmed (data not shown). **(C)** Volcano plot showing log2(fold change) of V, D, J gene frequencies between CD4+T and CD8+T cells (x-axis) and -log10(p-value) calculated by Cochran–Mantel–Haenszel test. Genes with most significant changes are annotated.

**Table S1.** Antibodies used for FACS staining

| Marker | Fluorochrome | Clone | Isotype | Supplier |
| --- | --- | --- | --- | --- |
| CD123 | BUV395 | 7G3 | Mouse IgG2a $\kappa$ | BD Biosciences |
| CD11c | APC-Cy7 | Bu15 | Mouse IgG1 $\kappa$ | Biolegend |
| CD14 | PE CF594 | MϕP9 | Mouse IgG2b $\kappa$ | BD Biosciences |
| CD137 | PE-Cy5 | 4B4-1 | Mouse IgG1 $\kappa$ | Biolegend |
| CD141 | PerCP-Cy5.5 | M80 | Mouse IgG1 $\kappa$ | Biolegend |
| CD19 | FITC | 4G7 | Mouse IgG1 $\kappa$ | BD Biosciences |
| CD20 | FITC | L27 | Mouse IgG1 $\kappa$ | BD Biosciences |
| CD3 | BV605 | SK7 | Mouse IgG1 $\kappa$ | Biolegend |
| CD4 | BV711 | RPA-T4 | Mouse IgG1 $\kappa$ | Biolegend |
| CD8A | AF700 | HIT8a | Mouse IgG1 $\kappa$ | Biolegend |
| CD8B | FITC | REA715 | Human IgG1 | Miltenyi Biotec |
| HLA-DR | BV785 | L243 | Mouse IgG2a $\kappa$ | Biolegend |
| EpCAM | Vioblue | HEA125 | Mouse IgG1 $\kappa$ | Miltenyi Biotec |
| CD45 | APC | HI30 | Mouse IgG1 $\kappa$ | BD Biosciences |
| CCR7 | PerCP-Cy5.5 | G043H7 | Mouse IgG2a $\kappa$ | Biolegend |
| CD56 | PE | NCAM16.2 | IgG2b, $\kappa$ | BD Biosciences |
| CD34 | PE-Cy7 | 581 | Mouse IgG1 $\kappa$ | Biolegend |
| CD3 | APC | SK7 | Mouse IgG1 $\kappa$ | Biolegend |
| THY1 | Af700 | 5E10 | Mouse IgG1 $\kappa$ | Biolegend |
| PEGFRa | PE | 16A1 | Mouse IgG1 $\kappa$ | Biolegend |
| PI16 | BV605 | RUO | Mouse IgG1 $\kappa$ | BD Biosciences |

**Table S2.** Probes used for smRNA FISH

| Gene ID | Cat. Number | Channel |
| --- | --- | --- |
| CSF2RA | 409341 | C1 |
| CCR7 | 410721 | C1 |
| LAMP3 | 468761-C2 | C2 |
| CD80 | 421471-C3 | C3 |
| CD8A | 560391-C3 | C3 |
| FOXP3 | 418471 | C1 |
| TNFRSF9 | 415171 | C1 |
| ITGAX | 419151 | C1 |
| FBN1 | 482478-C2 | C2 |
| COLEC11 | 542438 | C1 |
| ACTA2 | 311818-C3 | C3 |
| PDGFRA | 604488 | C1 |
| CDH5 | 437458-C3 | C3 |
| XCR1 | custom | C3 |
| FOXD4 | custom | C3 |
| GNG4 | custom | C2 |
| AIRE | custom | C1 |

**Table S3.**
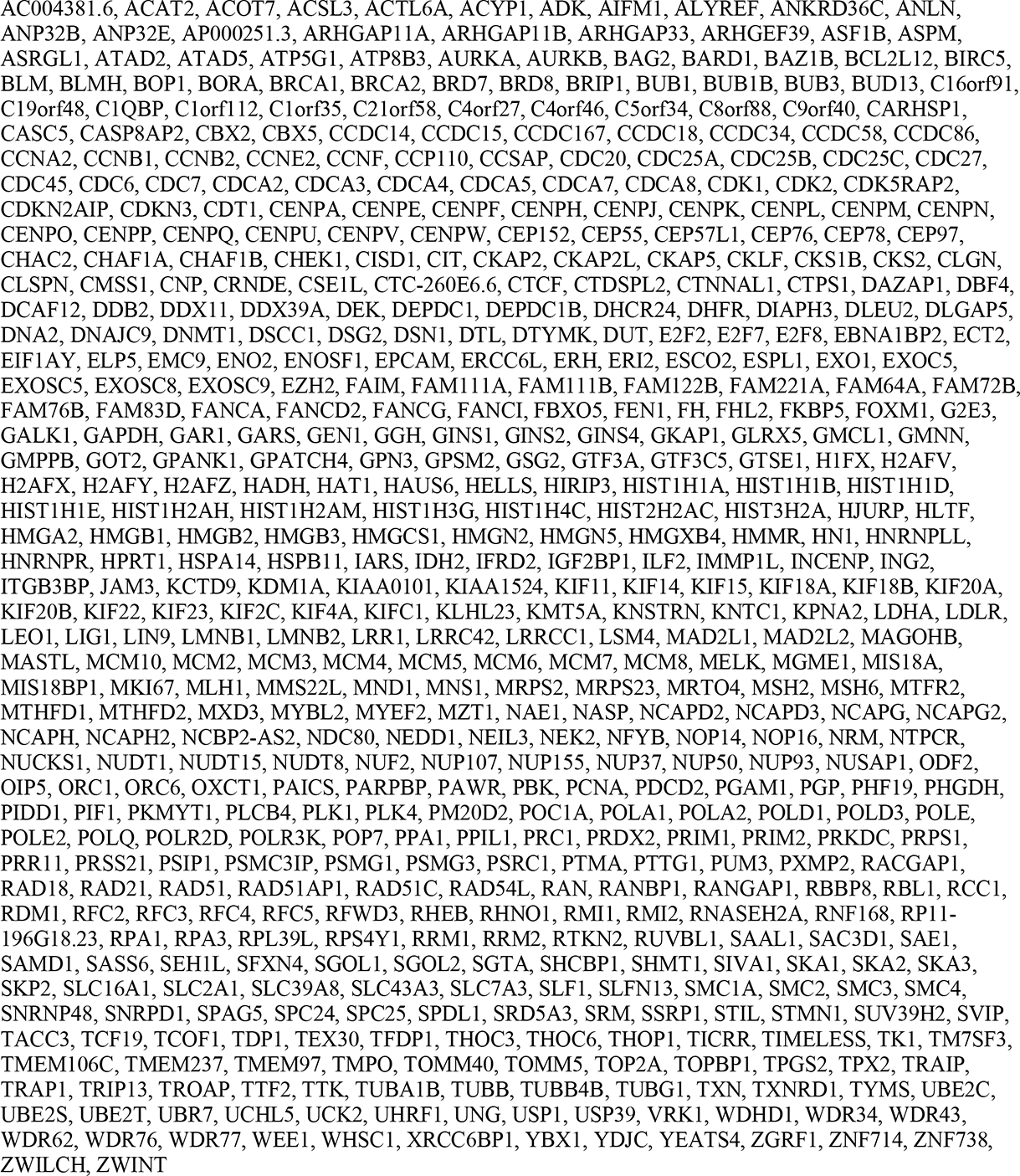
List of cell cycle genes (559 genes) defined and used in this study

